# Lateral surface pressure generated by nascent ribosomal RNA suppresses growth of fibrillar centers in the nucleolus

**DOI:** 10.1101/2021.09.09.459702

**Authors:** Tetsuya Yamamoto, Tomohiro Yamazaki, Kensuke Ninomiya, Tetsuro Hirose

## Abstract

Liquid-liquid phase separation (LLPS) has been thought to be the biophysical principle governing the assembly of the multiphase structures of nucleoli, the site of ribosomal biogenesis. Condensates assembled through LLPS increase their sizes to minimize the surface energy as far as their components are available. However, multiple microphases, fibrillar centers (FCs), dispersed in a nucleolus are stable and their sizes do not grow unless the transcription of pre-ribosomal RNA (pre-rRNA) is inhibited. To understand the mechanism of the suppression of the FC growth, we here construct a minimal theoretical model by taking into account the nascent pre-rRNAs tethered to the FC surfaces by RNA polymerase I. Our theory predicts that nascent pre-rRNAs generate the lateral osmotic pressure that counteracts the surface tension of the FCs and this suppresses the growth of the FCs over the stable size. The stable FC size decreases with increasing the transcription rate and decreasing the RNA processing rate. This prediction is supported by our experiments showing that RNA polymerase inhibitors increase the FC size in a dose-dependent manner. This theory may provide insight into the general mechanism of the size control of nuclear bodies.

**Significance statement:** The nucleolus, a site of pre-ribosomal RNA (pre-rRNA) production, has a characteristic multiphase structure, which has been thought to be assembled through liquid-liquid phase separation (LLPS). Although condensates assembled through LLPS grow by coarsening or coalescence as far as the components are available, the multiple inner phases, fibrillar centers (FCs), are dispersed in a nucleolus. To investigate the underlying mechanism, we constructed a minimal theoretical model by considering nascent pre-rRNAs tethered to RNA polymerase I at the FC surface. This model is supported by our experiments and explains previous experimental observations. This work shed light on the role of nascent RNAs to control the size of nuclear bodies.

## INTRODUCTION

In the interchromatin spaces, there are a variety of nuclear bodies, such as nucleoli (1-4), nuclear speckles (5), and paraspeckles (6). Many of the nuclear bodies are scaffolded by ribonucleoprotein (RNP) complexes, which are composed of RNA and RNA-binding proteins (RBPs). The class of RNAs, which are essential to the assembly of specific subcellular bodies, is called architectural RNA (arcRNA) (6-8). A growing number of researches suggest that some nuclear bodies are assembled via liquid-liquid phase separation (LLPS), which is driven by the multivalent interaction between RBPs bound to arcRNAs (6-8). Condensates produced by LLPS are spherical and increase their size by coarsening and/or coalescence due to the surface tension (macroscopic phase separation) (9).

Nucleoli are nuclear bodies, where ribosome assembly takes place (1,2). In the nucleoli, pre-ribosomal RNAs (pre-rRNAs) are transcribed by RNA polymerase I (Pol I) and maturated to construct ribosomes with ribosomal proteins. A nucleolus is not a uniform disordered liquid, but multiple microphases, called fibrillar centers (FCs), are dispersed in the sea of a granular component (GC) (Fig. 1A). Fibrillarins (FBLs), which are RBPs interacting with pre-rRNAs, are condensed to form another phase, called the dense fibrillar component (DFC), at the interfaces between FCs and the GC. The multiphase structures of nucleoli have been thought to be assembled via simple LLPS (10). However, this picture may not be complete because the FCs do not show coalescence or coarsening to increase their sizes. Indeed, when the Pol I transcription of pre-rRNA is inhibited, FCs show coalescence, as in the case of LLPS, and are excluded to the surface of the nucleolus (the excluded FCs are called nucleolar caps) (11). This implies that transcription somehow suppresses the growth of FCs. Ribosomal DNA (rDNA), from which pre-rRNAs are transcribed, is a repeat sequence of coding units (10 kb) intervened by the intergenic regions (30 kb). The transcriptionally active rDNA units and Pol I are localized at the surfaces of FCs where the transcription of pre-rRNAs takes place (3, 11-13). The DFC layer is probably assembled by the RNP complexes of nascent pre-rRNAs and the associated RBPs, such as FBLs (18). FBLs show phase separation with the physiological concentration and bind to nascent pre-rRNAs (10,13). These experimental results imply that nascent pre-rRNAs may act as ‘surfactants’ that suppress the growth of FCs.

**Figure 1.**
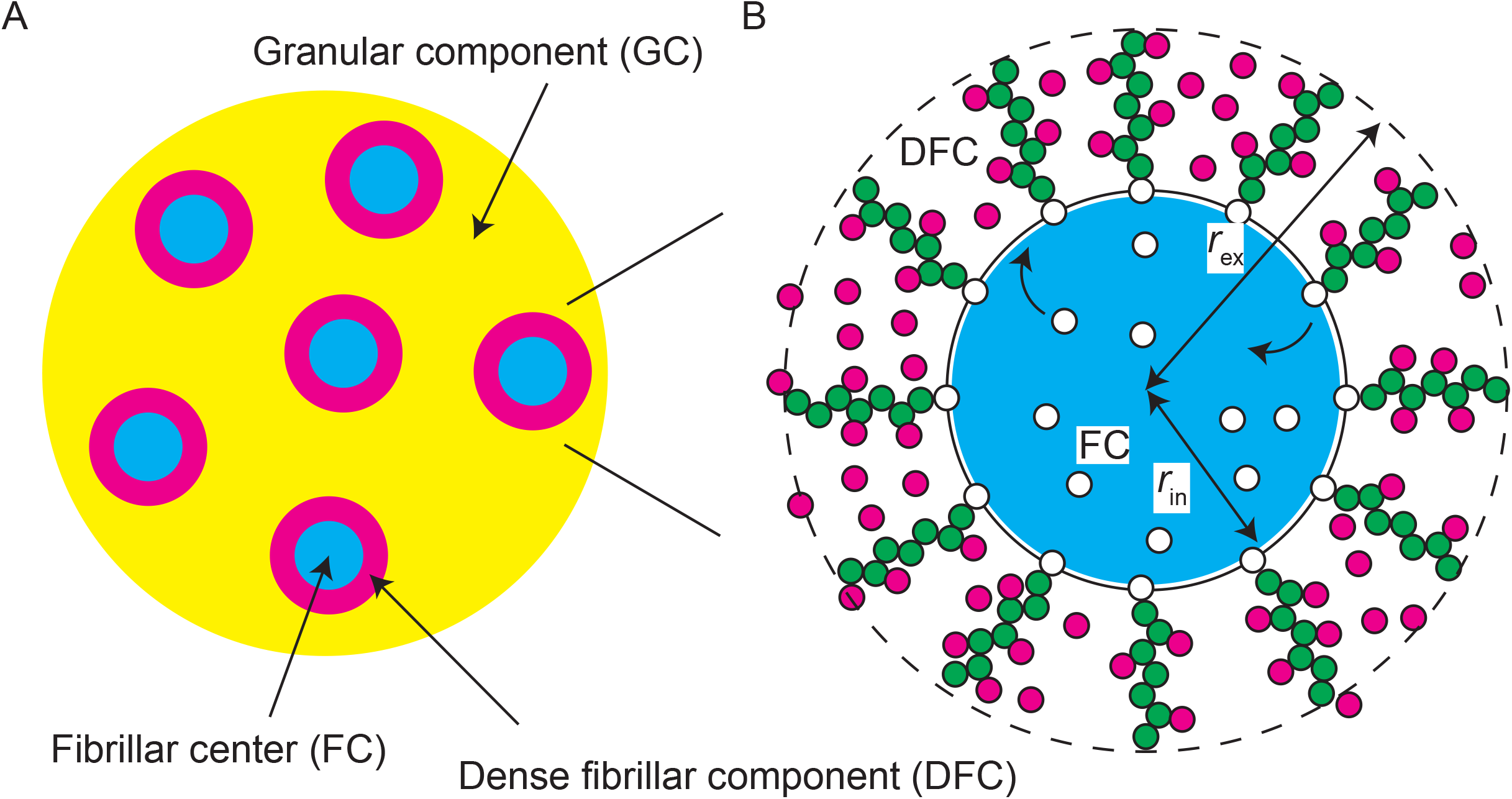
Multi-phase structure of a nucleolus. (A) A nucleolus is composed of multiple fibrillar center (FC) microphases in the sea of the granular component (GC). There is a layer of dense fibrillar component (DFC) between each FC and GC. (B) RNA polymerase I (Pol I) molecules (white particles) are entrapped in FC microphases (light blue) and the active rDNA units (black line) are localized at the surfaces of microphases. Nascent pre-rRNA transcripts (green particles) are thus at the surfaces of the microphase and form a DFC layer with RNA-binding proteins (magenta particles). The interface between FC and DFC is located at a distance *r*_in_ from the center and the interface between DFC and GC is located at a distance *r*_ex_.

Multiple paraspeckles scaffolded by NEAT1_2 arcRNAs are dispersed in a nucleoplasm (14). This situation is somewhat analogous to FCs in a nucleolus. Paraspeckles form the core-shell structure, where the two terminal regions of NEAT1_2 form the shell and the middle region of NEAT1_2 forms the core (15,16). We have recently shown that the terminal regions of NEAT1_2 in the shell suppresses the growth of paraspeckles, analogous to micelles of triblock copolymers (14,17). The size of paraspeckles increases with the NEAT1_2 transcription upregulation (14,18), which is opposite to the response of FCs in a nucleolus to transcription upregulation. This implies that the size control mechanism of FCs in a nucleolus is different from that of paraspeckles.

We here construct a simple theoretical model that predicts the contributions of nascent pre-rRNAs to the assembly of the multiphase structure of nucleolus. This model takes into account the phase separation, the transcription kinetics, and the interfacial effects, which are probably the minimum to account for the multiphase structure. Our theory predicts that the nascent pre-rRNAs are stretched to accommodate FBLs in the DFC layer and generate the lateral pressure that counteracts with the interfacial tensions. The size of FCs is determined by the balance of the interfacial tensions and the lateral pressure. Our theory predicts that the radius of FCs is proportional to the inverse of the square root of the transcription rate. The suppression of the FC growth by the pre-rRNA transcription results from the fact that these complexes are end-grafted to the FC surfaces via Pol I, in contrast to other condensates in which arcRNAs diffuse freely and increase their sizes by the transcription upregulation. To test our prediction, we experimentally measured the FC sizes and the pre-rRNA levels while the transcription rate is changed by the dose of BMH-21 or CX-5461, which specifically inhibits the Pol I transcription. The scaling exponent predicted by our theory is consistent with our experimental results, implying that the lateral pressure generated by nascent pre-rRNAs are possible mechanism of the size control of FCs. We anticipate that this theory can be a base theory to further look into the contribution of the multiphase structure in the function of nucleoli and can be extended to study the mechanism of the size control and the functions of other nuclear bodies, such as nuclear speckles and transcriptional condensates.

## RESULTS

### Model of nucleolus

We here construct a minimal model of a nucleolus to predict the size of FCs in the steady state. The nucleolus is composed of a GC in which multiple spherical FCs are dispersed (Fig. 1A). Pol I molecules (shown by white particles in Fig. 1B) are entrapped in the FCs (shown by a light blue droplet in Fig. 1B). The transcriptionally active repeat units of rDNA (shown by the black line in Fig. 1B) are localized at the surfaces of the FCs. We assume that the number of Pol I molecules *N*_pol_ and the number of transcriptionally active rDNA units *N*_act_ in the nucleolus are constant. Nascent pre-rRNAs (shown by chains of green particles in Fig. 1B) are localized at the surfaces of FCs and form a DFC layer (Fig. 1B). Recent experiment revealed that only the 5’ terminal external transcribed spacer (ETS) regions of nascent pre-rRNAs are localized at the DFC layer and that, among the proteins localized in the DFC layer, FBLs contributed most significantly to the localization of the terminal regions of nascent pre-rRNAs in the DFC layer (13). Motivated by this result, we take into account only the FBL-binding terminal regions of nascent pre-rRNAs as arcRNAs that scaffolds the DFC layers and only FBLs as RBPs that bind to these regions (see also the Discussion section). Recent experiments also revealed that the upstream half of ETSs are localized at the DFC layer and the downstream half is localized at the FC (13). This motivated us to assume that the terminal regions are spanning from the top to the bottom of the DFC layer. Some RBPs bind to the nascent pre-rRNAs and the others diffuse freely in the DFC layer (shown by magenta particles in Fig. 1B). The interface between FC and DFC is located at a distance *r*_in_ from the center and the interface between DFC and GC is located at a distance *r*_ex_ from the center (see Fig. 1B). For simplicity, we assume that all the FCs have equal volume and the sum of the volumes of microphases is fixed to *V*_m_. This assumption does not seem so far off although it has not been quantified (19).

The number of nascent pre-rRNAs at the surfaces of FCs is determined by the kinetics of transcription and RNA processing. We treat the transcription dynamics of pre-rRNA by using an extension of the model used by Stasevich and coworkers (20) (Fig. 2). Leaving the kinetic equations to Materials and Methods, we here discuss the processes of transcription and RNA processing that are taken into account in our model, see eqs. 6-10. Pol I in an FC binds to the transcription start site (TSS) of a transcriptionally active rDNA unit. The Pol I bound to the TSS starts transcription (Fig. 2A) or returns to the FC without starting transcription (Fig. 2A). The rate with which the Pol I bound to the TSS starts transcription is smaller than the rate with which this Pol I returns to the FC without starting transcription. The Pol I polymerizes a pre-rRNA while it moves uni-directionally towards the transcription termination site (TTS). The nascent pre-rRNAs are subject to co-transcriptional RNA processing (Fig. 2B). The FBL-binding terminal region is located at the ETS of pre-rRNA and is cleaved co-transcriptionally by endoribonuclease. The average time between the transcription initiation and the cleavage of the FBL-binding region is *τ*_pr_, where this time includes the elongation time to the cleavage site and the time necessary for the enzymatic reaction, see Fig. 2B and the Discussion section. Pol I reaches TTS at the average elongation time *τ*_e_ and is then released to the FC (*τ*_pr<_ *τ*_e_), see Fig. 2C. This model is designed to focus on the transcription of the FBLs of nascent pre-rRNAs and to treat the rest of the process as simple as possible. The kinetic equations of the Pol I transcription and co-transcriptional processing are derived by taking into account the above-mentioned processes, see Fig. 2 and eqs. 6-8. By solving these kinetic equations for the steady state, we found that the surface density *σ*_in_ is proportional to the radius *r*_in_ of FCs, see eq. 11 in Materials and Methods. We thus introduce a dimensionless parameter

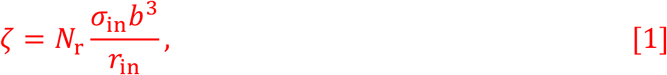

which is proportional to the transcription rate and processing time *τ*_pr_, while it is independent of the radius *r*_in_.

**Figure 2.**
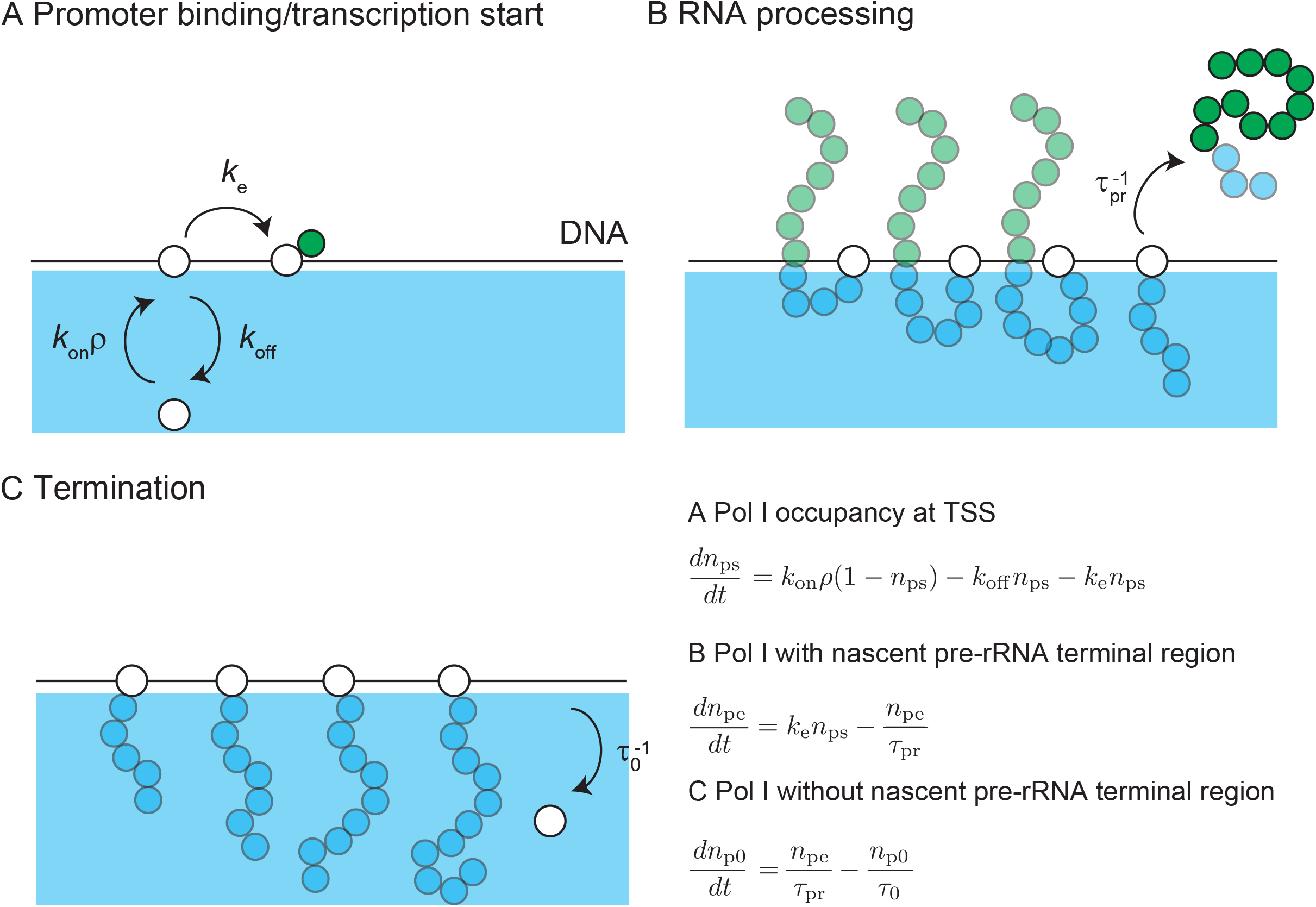
Model of transcription dynamics. DNA (black solid line) is localized at the surface of an FC (cyan). (A) RNA polymerase I (Pol I) in a microphase binds to the transcription starting site (TSS) of an active rDNA repeat unit. The bound Pol I starts transcription with the rate *k*e or returns to the microphase without starting transcription. (B) During the transcription, Pol I migrates uni-directionally towards the transcription terminating site (TTS) while polymerizing a nascent pre-rRNA. The terminal region of the nascent pre-ribosomal RNA (pre-rRNA), to which fibrillarin binds, are cleaved by the co-transcriptional RNA processing with the rate τ_pr_^-1^. The terminal region is released to GC. (C) After the cleavage of the terminal region, Pol I continues transcription until it reaches the TTS. At the TTS, Pol I is releaed to the FC with the rate τ_0_^-1^.

The stability of the system is quantified by the free energy. The free energy *F*_d_ of a DFC layer has 4 contributions: 1) the elastic free energy *f*_ela_ of nascent pre-rRNA, 2) the mixing free energy *f*_mix_ of RBPs and solvent, 3) the interaction free energy *f*_int_ between RBPs (both freely diffusing in the DFC layer and bound to nascent pre-rRNA), and 4) the binding free energy *f*_bnd_ of RBPs to the pre-rRNA terminal regions. The expressions of these free energy contributions are shown in eqs. 12–17 in Materials and Methods. The elastic free energy *f*_ela_ of flexible polymer chains, such as nascent pre-rRNAs, increases as the chains stretch because the number of possible conformations decreases (9). The thermal fluctuation mixes the components (nascent pre-rRNA units, RBPs, and solvent) and this contribution is quantified as the mixing free energy *f*_mix_. The interaction free energy *f*_int_ is the free energy contribution of the multivalent interactions between RBPs. The binding free energy *f*_bnd_ represents the free energy gain due to the binding of RBPs to nascent pre-rRNA units and the thermal fluctuations that dissociate RBPs from nascent pre-rRNA units. The free energy *F*_d_ takes into account the binding of RBPs to the terminal regions of nascent pre-rRNAs and the spherical geometry of the system in an extension of the Alexander model of polymer brush (21).

The free energy *F* of the system has the form

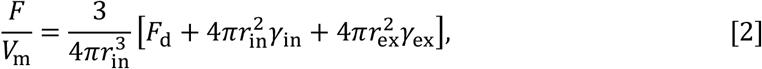

where *γ*_in_ is the interfacial energy per unit area (interfacial tension) at the interface between FC and DFC (*r* = *r*_in_) and *γ*_ex_ is the surface energy per unit area (interfacial tension) at the interface between DFC and GC (*r* = *r*_ex_). The external radius *r*_ex_ is determined by the conservation of the number of RNA units in the DFC layer in the steady state, see eq. 11 in the Materials and Methods. The interfacial tension results from the interactions between the components of interfacing phases. We therefore assume that the surface tensions, *γ*_in_ and *γ*_ex_, are proportional the local volume fraction of RBPs at the interfaces, with the proportional coefficient *γ*_p_ (see eqs. 18 and 19 in the Materials and Methods). The free energy *F*_d_ is a functional of the occupancy *α*_p_ of the pre-rRNA terminal regions by RBPs, the volume fraction *ϕ*_p_ of RBPs freely diffusing in the DFC layer, and the volume fraction *ϕ*_r_ of the FBL units, where these are functions of the distance *r* from the center of the FC.

In our theory, we take into account the non-equilibrium nature of the system via nascent pre-rRNAs produced at the surfaces of FCs by Pol I transcription. One may also think that the continuous addition of RNA units to nascent pre-rRNAs makes the conformations of their terminal regions different from those at the equilibrium. In a simple estimate, the local equilibrium approximation that uses the equilibrium chain conformations is effective in the length scales smaller than the diffusion length 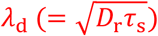, where *D*_r_ is the diffusion constant of a pre-rRNA unit and *τ*_s_ is the elongation time to produce a pre-rRNA unit. The diffusion length *λ*_d_ is estimated as ∼ 3 μm (see Table 1) and is larger than the typical thickness, ∼ 200 nm, of the DFC layer: the local equilibrium approximation is effective to treat the conformation of the pre-rRNAs in the DFC layer. The occupancy *α*_p_, the volume fractions *ϕ*_p_ and *ϕ*_r_ of RBPs and nascent pre-rRNA, and the radius *r*_in_ thus turn over towards the minimum of the free energy, eq. 2.

**Table 1.** The values of the parameters used to estimate the diffusion length *λ*_d_. The diffusion constant of a RNA unit is estimated as 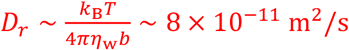 and the time to synthesize a RNA unit is estimated as *τ*_s_ = *b/υ*_e_ ∼ 0.12 s by using the values listed in this table. We estimated *b* from the persistence length of single-stranded DNA.

| Parameter | Meaning | Value | Ref. |
| --- | --- | --- | --- |
| $b$ | Length of a RNA unit | 4 nm ≈ 12 b | 22 |
| $\eta_w$ | Water viscosity | 1 mPa · s | |
| $v_e$ | Elongation rate | 6 kb/min | 23,24 |
| $T$ | Absolute Temperature | 300 K | |

### Parameter estimate suggests that the terminal regions of nascent pre-rRNAs are highly stretched

The independent parameters involved in our theory and their estimates are summarized in Table 2. The kinetic constants involved in the Pol I transcription are included in *ζ*, see eq. 1. The energy gain due to the binding of RBPs to the pre-rRNA terminal regions is represented by *εk*_B_*T*. We represent the magnitudes of the multivalent interaction between RBPs by using the Flory interaction parameter *χ*. We neglect the dependence of the magnitudes of the interaction between RBPs on their binding state and also assume that solvent molecules and naked pre-rRNA units are equivalent in terms of the magnitudes of the interaction. This highlights the fact that the multivalent interaction between FBLs drives phase separation *in vitro*, not to lose the essence by introducing many interaction parameters. The volume fraction *ϕ*_p0_ of RBPs in the nucleoplasm is taken into account via the chemical potential *µ*_p_ = *k*_B_*T* log *ϕ*_p0_.

**Table 2.** List of independent (dimensionless) parameters involved in our theory. Estimate of the unit length *b* of pre-rRNA and the absolute temperature are shown in Table 1. The surface density *σ*_in_ was estimated by the number of Pol I (50 Pol I per rDNA, 300 copies of active rDNA repeat per cell, ∼ 10 FCs per nucleolus, ∼ 2 nucleoli per cell) and the typical radius of FCs *r*_in_ ∼ 100 nm (3,13). The surface tension *γ*_p_ can be estimated as 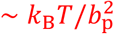 (25,26), where the size *b* of FBL is ≈ 4 nm (27). The number of units *N* in the FBL-binding region of pre-rRNA is estimated by using its length ∼ 400 b (13). The chemical potential of FBL was estimated by using its concentration ∼ 1 μM (10).

| Parameter | Meaning | Estimate | Value | Ref |
| --- | --- | --- | --- | --- |
| $\zeta$ | Rescaled pre-rRNA surface density | $< 0.12$ | Varied | 13 |
| $\frac{\gamma_p b^2}{N_r k_B T}$ | Rescaled surface tension per RBP | $\sim 0.03$ | 0.03 | |
| $\chi$ | Interaction parameter | $> 10$ | 12 | |
| $\epsilon$ | RBP-RNA binding energy | $> 10$ | -12 | |
| $\mu_p/(k_B T)$ | RBP chemical potential | -10 | -10 | 10 |

The interaction parameter *χ* and the binding energy *ε* are important quantities of polymer physics, but are not experimentally well characterized. In the limit of vanishing volume fraction of nascent RNAs, *σ*_in_ → 0, our model returns to the theory of the binary mixture of RBPs and solvent molecules (25), see eqs. 3 and 4. The latter theory predicts that the phase separation happens at the threshold interaction parameter *χ*_tr0_ (= −*µ*_p_/(*k*_B_*T*)). FBL molecules show phase separation *in vitro* with the physiological concentration (10), implying that *χ* > *χ*_tr0_. If FBLs bind to the pre-rRNA terminal regions even in a dilute solution, it implies that *ε* > −*µ*_p_/(*k*_B_*T*). The size of the FBL of the terminal region of a nascent pre-rRNA is ≈ 30 nm in the relaxed state (≈ 130 nm in the maximally stretched state), while the thickness of a DFC layer is ≈ 100 nm, implying that FBLs are highly stretched.

### The multivalent interaction between FBLs localizes these RBPs to the DFC layer and stretches the terminal regions of nascent pre-rRNAs

The occupancy *α*_p_, the volume fractions *ϕ*_p_ and *ϕ*_r_ of RBPs and nascent pre-rRNA at the free energy minimum is derived by using the variational principle, see sec. S1B in the SI Appendix. For cases in which the curvature of the surfaces of FCs is not negligible, *α*_p_, *ϕ*_p_, and *ϕ*_r_ are not uniform in the DFC layer. To avoid complexity arising from the geometry of the system, we first analyze the occupancy *α*_p_ and the volume fractions, *ϕ*_p_ and *ϕ*_r_, in a DFC layer on a planer surface. This is indeed the limit of large FCs and the essential feature of the assembly of DFC layers is included in this analysis. In the planer geometry, the chemical potential of RBPs and the osmotic pressure is uniform in the DFC layer.

The magnitude of the multivalent interactions between RBPs (either bound to the pre-rRNA terminal regions or freely diffusing) increases with increasing the interaction parameter *χ*. Although it is not straightforward to experimentally control the interaction parameter *χ* of FBLs, it is instructive to analyze the composition of the layer occupied by the FBLs of nascent pre-rRNAs as a function of the interaction parameter *χ*, see Fig. 3. For cases in which the interaction parameter *χ* is small (which corresponds to self-association defective FBL ΔIDR mutant (13)), the major component of the layer is solvent (nucleoplasm), see the cyan line in Fig. 3. For *χ*< *χ*_s_, the excluded volume interaction between nascent pre-rRNA units dominates the attractive interaction between RBPs bound to these nascent pre-rRNAs (good solvent regime). The interaction parameter *χ*_s_ at the crossover is 2 in the limit of *ε* ≪ *µ*_p_/(*k*_B_*T*), such as the case of Fig. 3, see also eq. S31 in the SI Appendix for the general expression. The inverse of the volume fraction *ϕ*_r_ of nascent pre-rRNA is proportional to the extent of the stretching of the terminal regions of nascent pre-rRNAs. The terminal regions of these RNAs are somewhat stretched for *χ* ≈ 0 due to the excluded volume interaction between the pre-rRNA units (good solvent regime). The terminal regions shrink as the magnitude *χ* of the attractive interaction between RBPs bound to the terminal regions of nascent pre-rRNAs, see the green line in Fig. 3. For *χ*_s<_ *χ*< −*µ*_p_/(*k*_B_*T*), the attractive interaction dominates the excluded volume interaction (poor solvent regime) and the terminal regions are collapsed with RBPs bound to these regions (melt regime), see the green line in Fig. 3.

**Figure 3.**
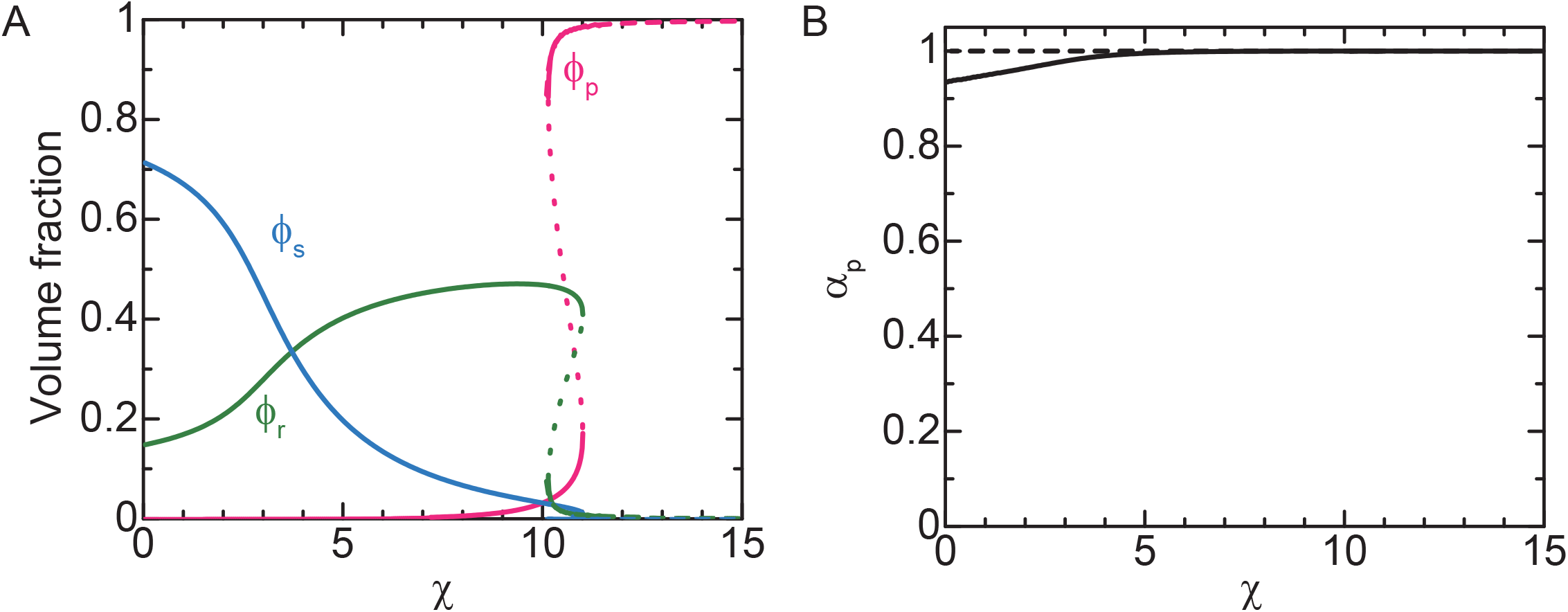
Composition of planer DFC layer vs interaction parameter χ. (A) The volume fraction φ_r_ of pre-rRNA units (green), the volume fraction φ_p_ of freely diffusing RBPs (magenta), the volume fraction φ_s_ of solvent molecules (cyan) in a DFC layer are shown as functions of the interaction parameter χ. (B) The occupancy α_p_ of RNA terminal region by RBPs is shown as a function of the interaction parameter χ. We used μ_p_/(*k*_B_*T*)=-10.0, ε=-12.0, σ_in_*b*^2^=0.05, and Π_ex_*b*^3^/(*k*_B_*T*)=0.0 for the calculations, see also Table 1. The solid lines are derived by numerically solving eqs. S15, S16, and S20 in the SI Appendix. The broken lines are derived by using eqs. S47 (shown for χ>10.1).

At a threshold interaction parameter, *χ*_th_ ≈ −*µ*_p_/(*k*_B_*T*), the volume fraction *ϕ*_p_ of freely diffusing RBPs jumps to a large value, see the magenta line in Fig. 3. In contrast, the volume fraction of the solvent (nucleosol) jumps to almost zero, see the cyan line in Fig. 3. The feature of the layer for *χ* > *χ*_th_ is analogous to the DFC layer observed experimentally (which we thus call DFC regime). The volume fraction *ϕ*_r_ of nascent pre-rRNA units jumps to a small value at *χ* ≈ *χ*_th_, see the green line in Fig. 3, implying that the terminal regions of the nascent pre-rRNAs are stretched at the transition. Because the volume fraction *ϕ*_r_ is smaller than the case of *χ* = 0, the stretching is not solely due to the fact that RBPs in the DFC layer behave as an athermal solvent to the RNP complexes of RBPs and the pre-rRNA terminal regions. The terminal regions stretch so that the DFC layer can accommodate more RBPs to decrease the free energy due to the multivalent interactions between them. The stretching thus may be a transport mechanism of the ETS of pre-rRNAs towards the DFC layer.

### The lateral pressure generated by nascent pre-rRNAs suppresses the growth of FCs

Now we discuss the DFC layers at the surfaces of spherical FCs to derive the radius of FCs in the steady state. The first (functional) derivatives of the free energy with respect to *α*_p_, *ϕ*_p_, and *ϕ*_r_ are zero in the steady state. In the spherical geometry, these conditions correspond to the fact that the chemical potential of RBPs and the chemical potential of pre-rRNA units, instead of the osmotic pressure, are uniform, see sec. S1B for the mathematics of the free energy minimization. We treat the DFC regime in which the volume fraction of solvent molecules is negligibly small and the occupancy *α*_p_ of the pre-rRNA terminal regions is approximately unity, see Table 2 and Fig. 3. We performed numerical and analytical calculations within these conditions, see sec. S3A in the SI Appendix.

In the spherical geometry, the volume fraction *ϕ*_r_ of pre-rRNA units is not uniform in the DFC layer and decreases with the distance *r* from the center (*r*_in_ < *r*< *r*_ex_). It is because the elastic stress (proportional to the number of pre-rRNAs per unit area) generated by pre-rRNA decreases with increasing *r*, see the solid green line in Fig. 4. We performed the asymptotic analysis for *ϕ*_r_ ≪ 1 by expanding the chemical potentials of proteins and pre-rRNA units in a power series and neglected higher order term of *ϕ*_r_. This leads to an asymptotic expression, 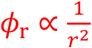, which reasonably explains the numerical calculation, see the solid dark green line and broken light green line in Fig. 4 and also eq. S55 in the SI Appendix for the explicit form. The rest of the space in the DFC layer is filled by RBPs, see the inset of Fig. 4.

**Figure 4.**
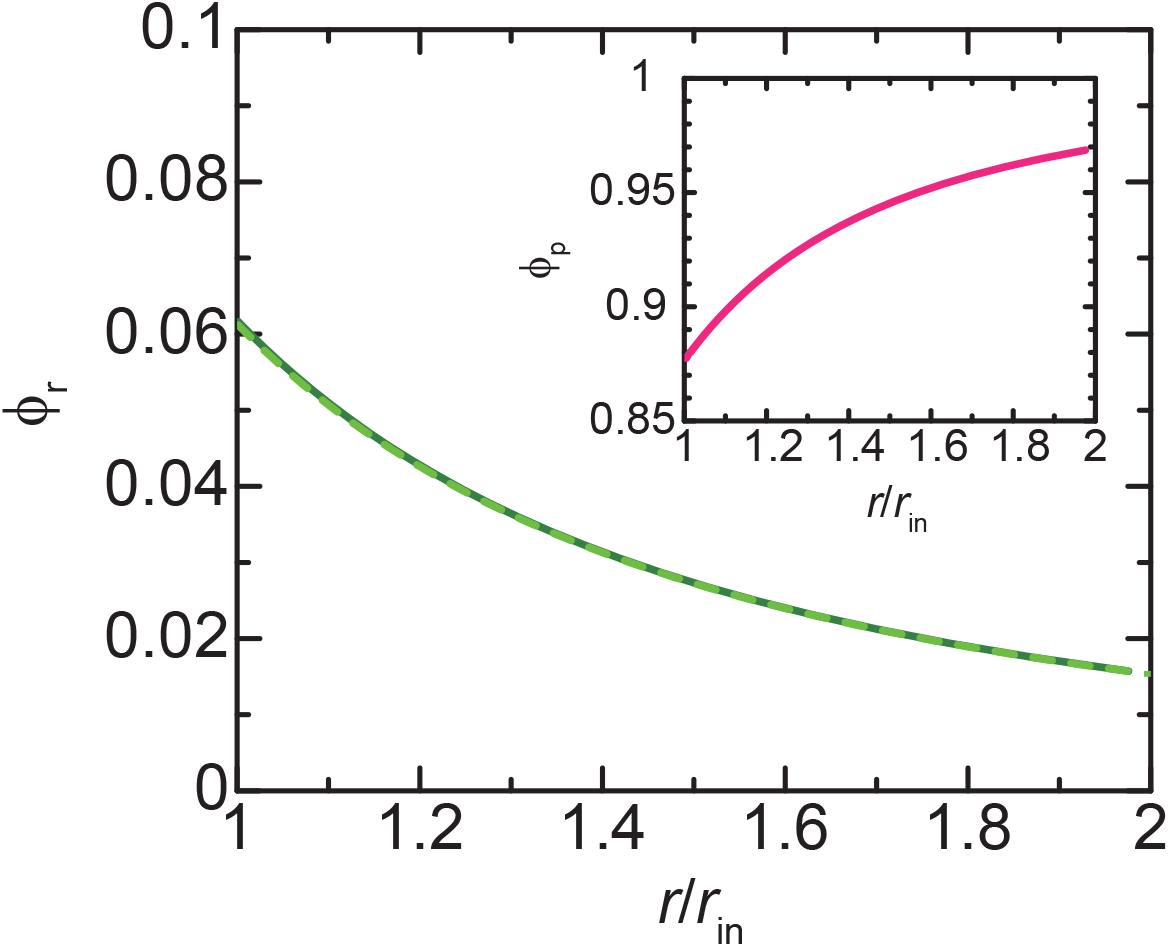
Profile of volume fraction φ_r_ of pre-rRNA units in DFC layer. The volume fraction of nascent pre-rRNA units is shown as a function of the position r in the DFC layer (*r*_in_<*r*<*r*_ex_). The solid dark green line is derived by numerically calculating eqs. S13, S15, and S16 in the SI Appendix for ζ=0.06 with the condition that the volume fraction of solvent is zero and the occupancy of the terminal regions of pre-rRNAs by RBPs is unity. The values of other parameters are summarized in Table 1. The broken light green line is derived by using eq. S59 in the SI Appendix with φ_ex_=0.0157 and *r*_ex_=1.98 (which were derived from the numerical calculation to obtain the light green line). The volume fraction of freely diffusing RBPs is shown in the inset.

We used the profile of the pre-rRNA units to derive the free energy as a function of the radius *r*_in_ of FCs. For cases in which the Pol I transcription is inhibited, the free energy of the system is a monotonically decreasing function of the radius *r*_in_, implying that, the radius of FCs increases with time by coarsening or coalescence. For cases in which the Pol I transcription is active, the free energy generated by the terminal regions of the nascent pre-rRNA is an intriguing function of *r*_in_, which has a minimum, see the solid black line in Fig. 5A. Indeed, the surface free energy, the second and third terms of eq. 2, is not significant for the values of parameters estimated in Table 2, see the broken cyan line in Fig. 5A. In such case, the free energy is simpler if it is shown as a function of the ratio *r*_ex_/*r*_in_, see Fig. 5B and eq. S59 in the SI Appendix.

**Figure 5.**
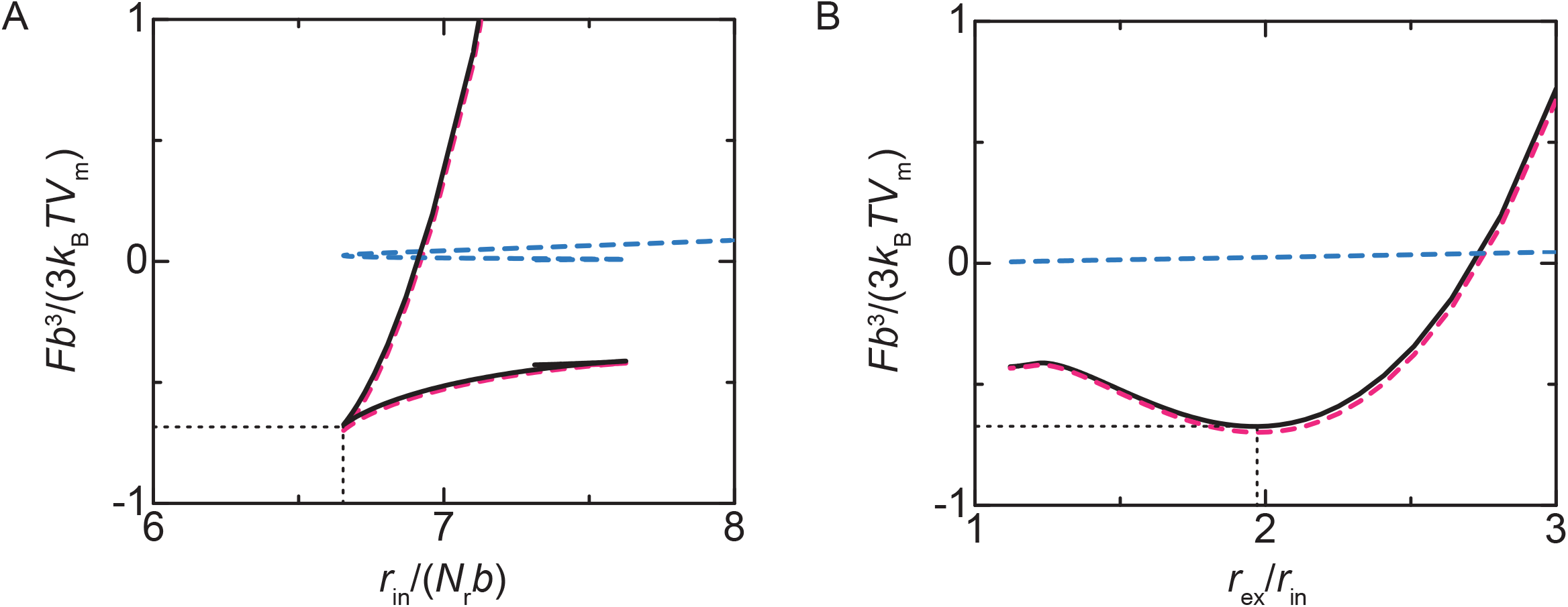
Free energy *F* of nucleolus. The free energy *F* of the system is shown as a function of the radius *r*_in_ of FCs (a) and the ratio *r*_ex_/*r*_in_ of the external radius to the internal radius (b). The black solid line is the total free energy, including the free energy of DFC layers (shown by the magenta broken line, the first term of eq. 2) and the surface free energy (shown by the cyan broken line, the second and third terms of eq. 2) for ζ=0.06. The values of other parameters are summarized in Table 1.

The first derivative of the free energy with respect to the radius *r*_in_ is zero at the minimum of the free energy,

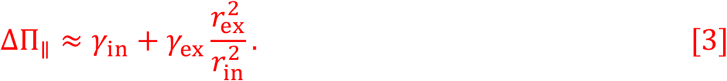

Eq. 3 represents the surface tensions, *γ*_in_ and *γ*_ex_, are balanced to the lateral osmotic pressure Δ∏_∥_ generated in the DFC layer (the complete form is shown in eq. 21). The radius *r*_in_ at the minimum of the free energy decreases with increasing the transcription level, see Fig. 6. The radius *r*_in_ at the minimum of the free energy has an asymptotic form

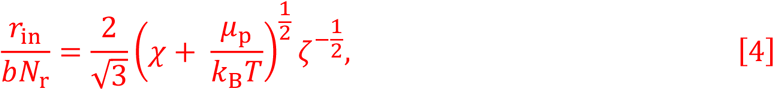

see also the broken line in Fig. 6. Eq. 4 is derived by expanding the free energy in a power series of *ϕ*_r_ and by using the asymptotic form of *ϕ*_r_, see sec. S3A in the SI Appendix for the details of the derivation. We also neglected the surface free energy, the second and third terms of eq. 2, because it is not significant with our estimate in Table 2. Our theory therefore predicts that the radius of FCs decreases with the inverse of the square root of the transcription rate. At the minimum of the free energy, the ratio of the radii is 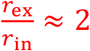, which roughly agrees with the fact that the radius of FCs and the thickness of DFC layers are the same order of length scales, ∝ 100 nm, see Fig. 5B.

**Figure 6.**
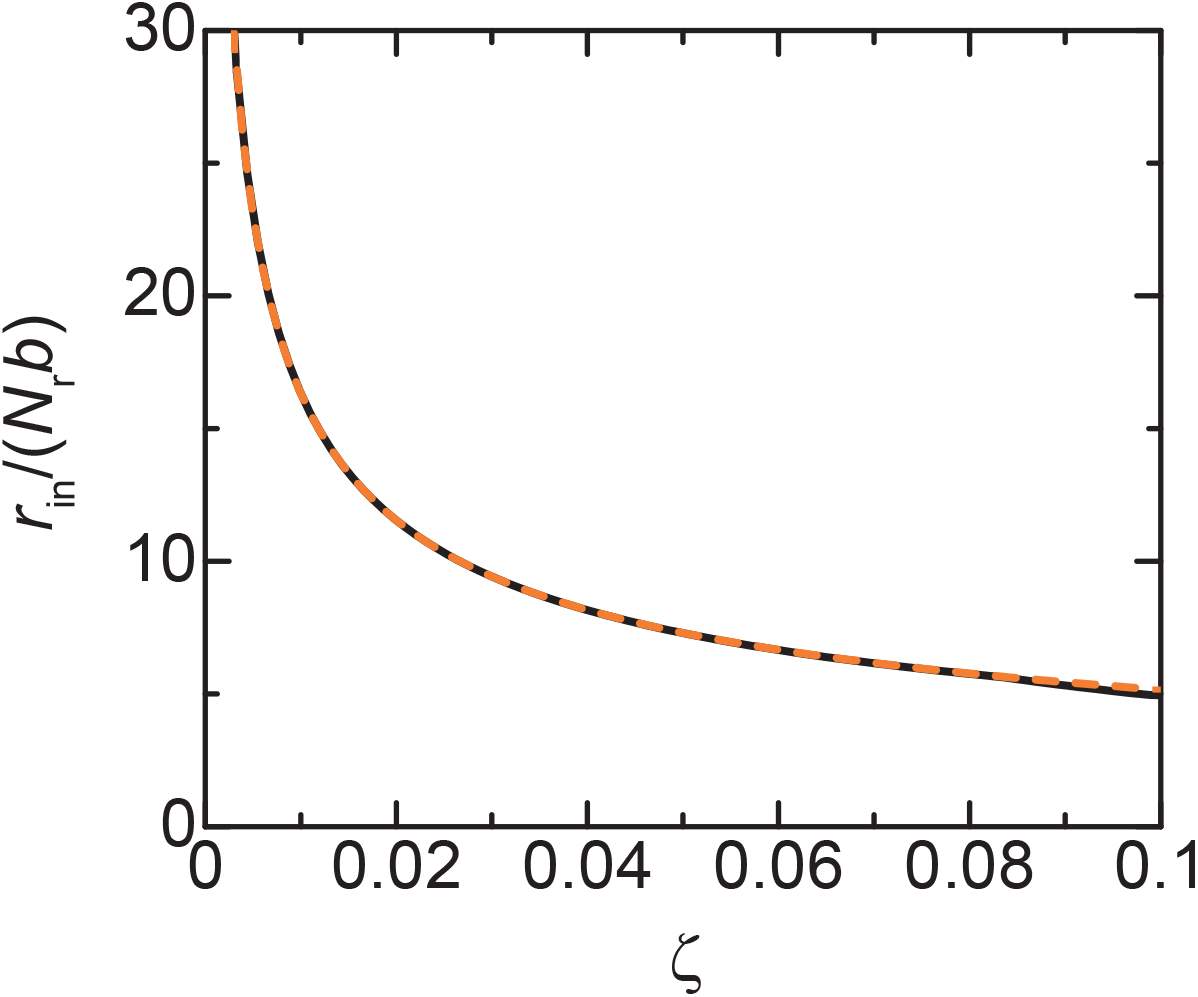
Radius *r*_in_ of FCs vs rescaled transcription rate ζ. The radius *r*_in_ of FCs at the free energy minimum is shown as a function of rescaled transcription rate ζ. The solid dark green line is derived by numerically calculating eqs. S13, S15, and S16 in the SI Appendix with the condition that the volume fraction of solvent is zero and the occupancy of the terminal regions of pre-rRNAs by RBPs is unity. The orange broken line is derived by using eq. 5. The parameters used for the calculations are summarized in Table 1.

### Mild inhibition of Pol I increases the size of FCs

The main prediction from our theoretical model is that the size of FCs increases by reducing the expression level of nascent pre-rRNAs. To experimentally test this prediction, we used BMH-21 and CX-5461, specific Pol I inhibitors, which reduce nascent pre-rRNA levels in a dose-dependent manner (28,29) (Fig. S1A and B). In untreated cells, small foci of UBF (a marker for FCs) and FBL (a marker for DFC) proteins were dispersed within the nucleoli as previously reported (11) (Figs. 7A, S2A, S2B, and S3A). In contrast, UBF and FBL proteins were relocalized to nucleolar caps in the presence of high doses of BMH-21 (0.5 µM) or CX-5461 (2 µM), as reported (11,28) (Figs. 7A, S2A, S2B, and S3A). Strikingly, the FCs were larger in cells treated with the medium doses of BMH-21 (0.0625, 0.125, and 0.25 µM) or CX-5461 (0.25, 0.5, and 1 µM) than in untreated cells (Figs. 7A, S2A, S2B, and S3A). We then quantified the size of the FCs under these untreated and mildly treated conditions (Figs. 7B, 7C, S3B, S3C, and S4). Indeed, the longest axis (Lx) and area of the FCs increase with increasing the dose of BMH-21 and CX-5461. As theoretically predicted, the Lx and area of the FCs increase according to pre-rRNA level reduction (Figs. 7D, 7E, S3D, and S3E).

**Figure 7.**
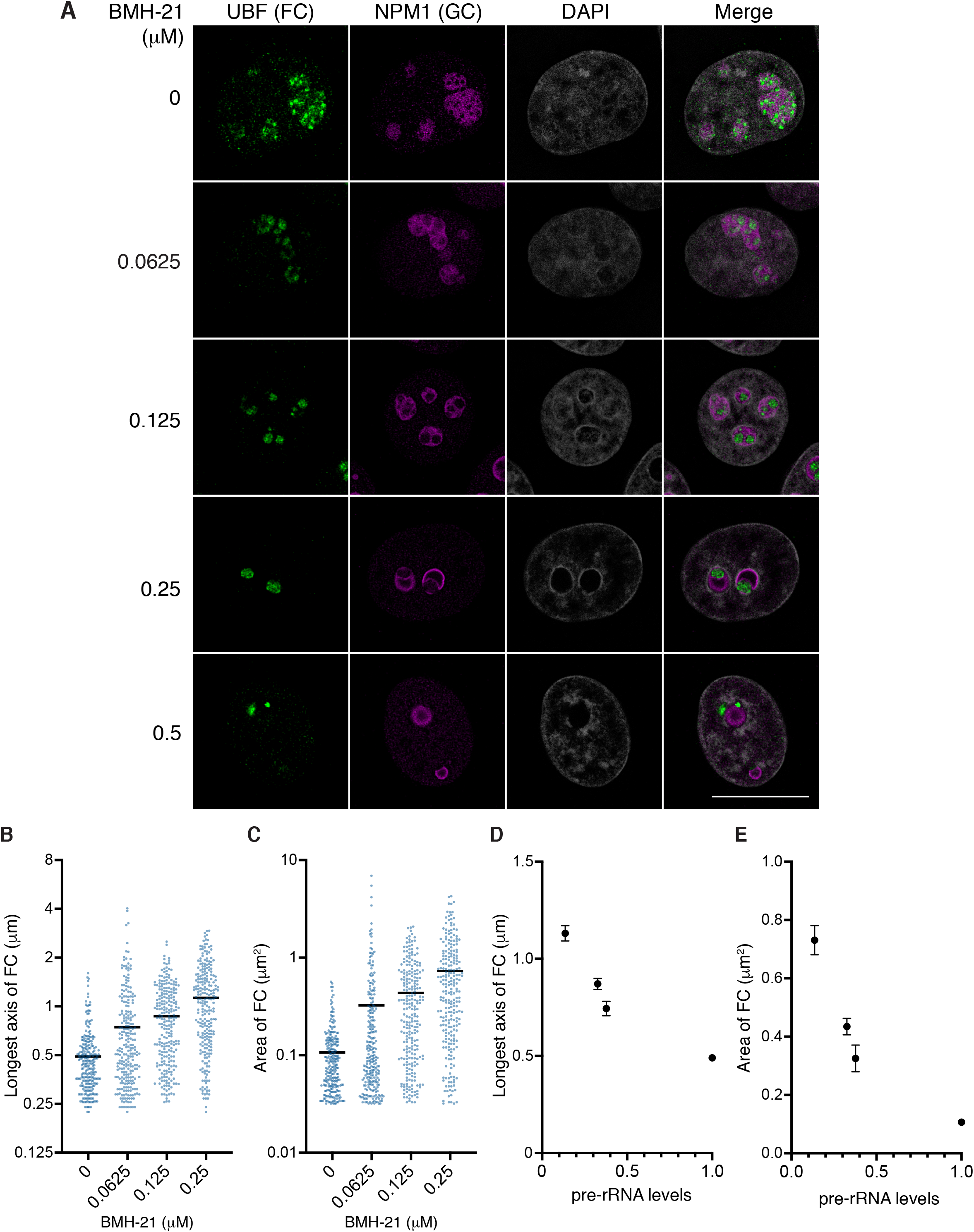
Mild Pol I inhibition by BMH-21 increases the size of FCs. (A) Immunofluorescence of UBF (FC) and NPM1 (GC) in HeLa cells with or without BMH-21 treatments. Scale bar, 10 μm. (B and C) Quantification of longest axis (B) and area (C) of the FCs in cells under indicated conditions. Each scatter dot plot shows the mean (black line). Dots indicate all points of quantified data (n = 250). Mean longest axes of the FCs is shown below: 0 μM: 0.4907 μm, 0.0625 μM: 0.7440 μm, 0.125 μM: 0.8710 μm, 0.25 μM: 1.131 μm. Mean areas of the FCs are shown below: 0 μM: 0.170 μm^2^, 0.0625 μM: 0.3255 μm^2^, 0.125 μM: 0.4347 μm^2^, 0.25 μM: 0.7312 μm^2^. Statistical analyses using Kruskal-Wallistest with Dunn’s multiple comparison test were performed and the results are shown as follows. (B) 0 μM vs 0.0625 μM: P < 0.0001, 0 μM vs 0.125 μM: P < 0.0001, 0 μM vs 0.25 μM: P < 0.0001, 0.0625 μM vs 0.125 μM: P < 0.0001, 0.0625 μM vs 0.25 μM: P < 0.0001, 0.125 μM vs 0.25 μM: P = 0.0008. (C) 0 μM vs 0.0625 μM: P = 0.0003, 0 μM vs 0.125 μM: P < 0.0001, 0 μM vs 0.25 μM: P < 0.0001, 0.0625 μM vs 0.125 μM: P < 0.0001, 0.0625 μM vs 0.25 μM: P < 0.0001, 0.125 μM vs 0.25 μM: P = 0.0010. (D and E) Graphs showing the mean longest axis (D) and area (E) of the FCs with SEM vs pre-rRNA expression levels. The pre-rRNA expression level in untreated cells is defined as 1.

### FC radius is a power function of the pre-rRNA transcription level with an exponent of approximately -0.5

To compare our theory and experiments more quantitatively, we curve-fit the square root of the area of FCs as a function of the transcription level by a power function (Fig. 8). The scaling exponent that describes the dependence of the square root of the area of FCs on the transcription level is a universal quantity that does not depend on specific values of the parameters. Checking such scaling exponents with experiments is the first thing to do to test theories in polymer physics (30). By fitting the data with a power function, we found that the scaling exponent is -0.49 for the case of the BMH-21 treatment, -0.46 for the case of the CX-5461, and -0.48 if both data are fitted. The scaling exponent predicted by our theory -0.5 reasonably agrees with our experiments, see eq. 4.

**Figure 8.**
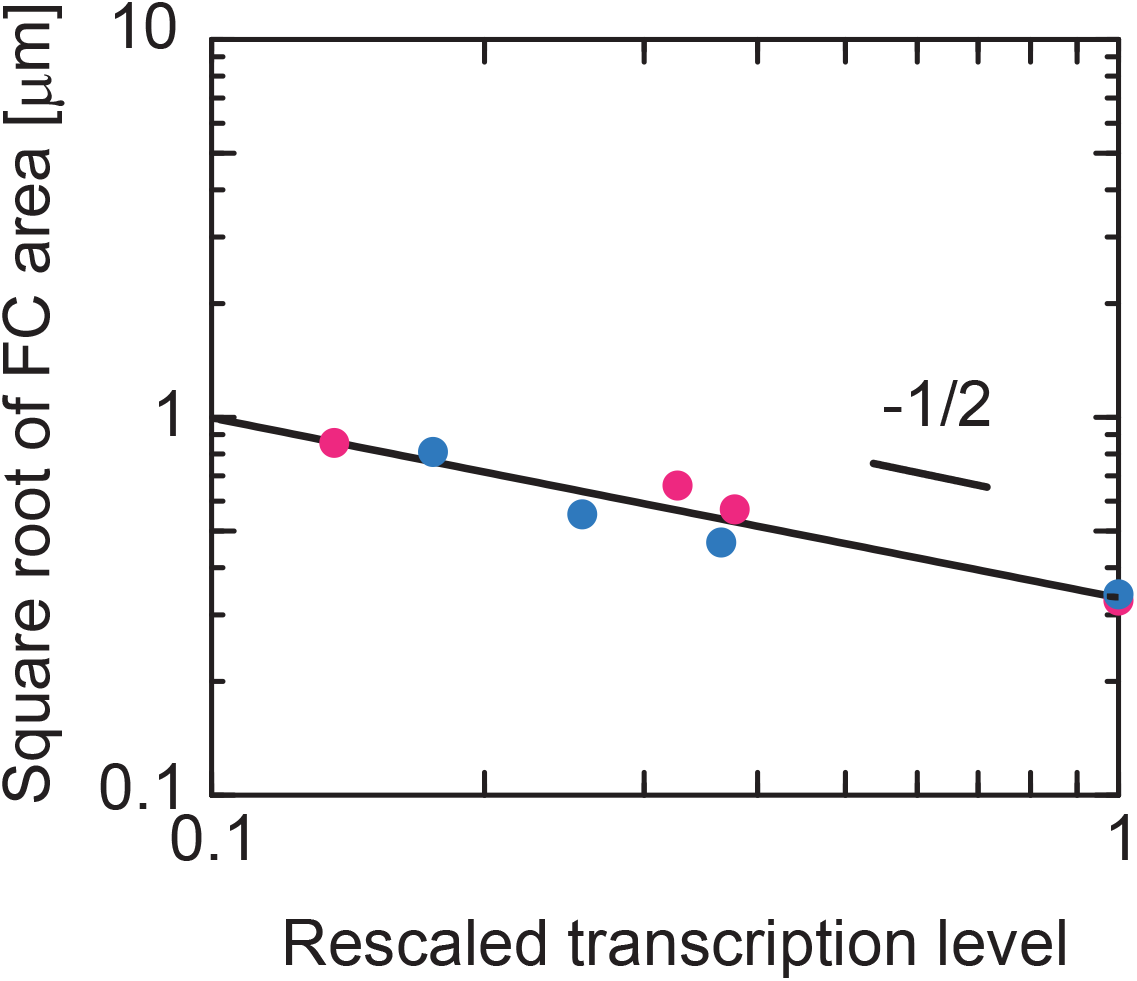
Exponent that account for the dependence of the size of FCs on pre-rRNA expression level. The data in Fig. 7E was shown in the double-logarithm plot and fitted with a power function. The magenta and cyan dots are the results by suppressing the Pol I transcription by using BMH-21 and CX-5461. The slope of the double-logarithm plot is the exponent that accounts for the dependence of the radius of FCs on the transcription level. The curve fitting shows that the exponent is -0.49 for the case of the BMH-21 treatment, -0.46 for the case of the CX-5461 treatment, and -0.48 if both data are fitted.

**Figure 9.**
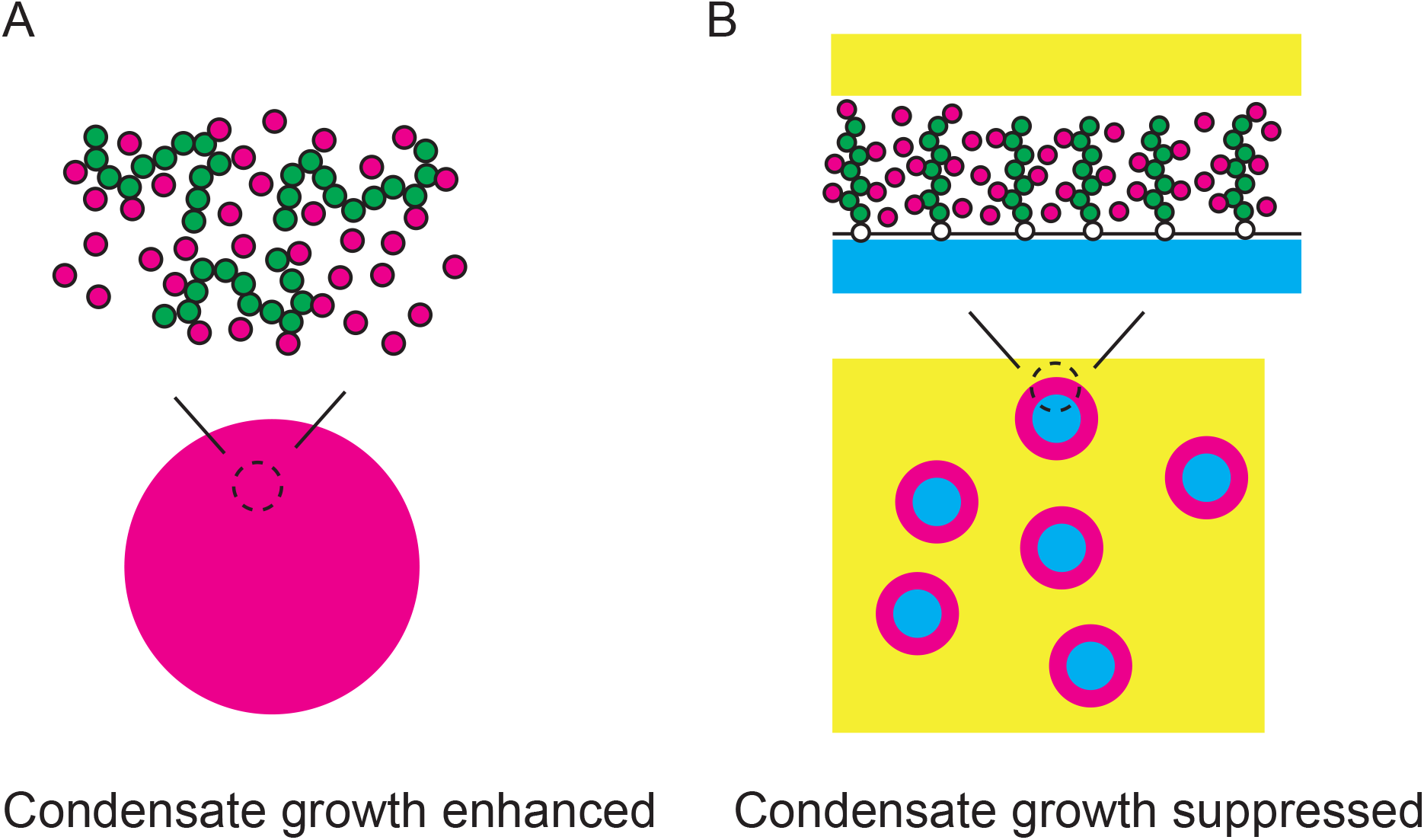
Summary of results. RNP complexes of RNAs and RBPs enhance or suppress the growth of condensates depending on whether the RNP complexes are mobile in the interior or tethered to the surfaces of the condensates. A. The multi-valent interaction between the RNP complexes enhance the growth of the condensates if these condensates are assembled by the RNP complexes. B. The multi-valent interaction between the RNP complexes suppress the growth of the condensates if these complexes are tethered to the surfaces of the condensates assembled by other RNAs and proteins.

## DISCUSSION

Our theory predicts that the terminal regions of nascent pre-rRNAs at the surfaces of FC microphases in a nucleolus generate the lateral osmotic pressure and suppress the growth of microphases. Our theory predicts that the radius of FCs is proportional to the inverse of the square root of the transcription rate. This prediction is consistent with our experiments. The parameter *ζ* is also proportional to the inverse of the rate 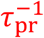 of co-transcriptional processing that cleaves the terminal regions of pre-rRNA. Our theory thus predicts that the radius of FCs is proportional to the square root of the processing rate 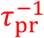. The inhibition of processing induced by ribosomal protein depletion, such as uL18 (RPL5) and uL5 (RPL11), changes the multiphase structure of nucleoli (31). Indeed, recent experiments have demonstrated that long pre-rRNAs are accumulated in RPL5 knockdown cells and the size of FCs in such cells is smaller than that in the wild type (32). The consistency between our theory and experiments suggests that the growth of FCs is suppressed by the terminal regions of nascent pre-rRNAs localized at the surfaces of FCs. This implies that the elastic energy of the terminal region is the driving force of the transport of the 5’ ETSs cleaved by RNA processing towards GC.

In many cases, the multivalent interaction between complexes of RNA and RBPs drives the growth of condensates (33,34) (Fig. 8A). The size of such condensates increases with increasing the transcription rate (14,17,35). In contrast, for the case of FCs in nucleoli, the multivalent interaction between the complexes of nascent RNA and RBPs rather suppresses the growth of FCs (11). The size of FCs indeed decreases with increasing the transcription rate. Our theory predicts that the suppression of the growth of FCs by transcription results from the fact that pre-rRNAs are end-grafted to the surfaces of FCs via Pol I (Fig. 8B).

Our theory predicts that the function of the terminal regions of nascent pre-rRNAs is analogous to surfactants, such as lipids. If the surfactants at an interface between two immiscible liquids are dense enough to form a monomolecular liquid film, these surfactants decrease the interfacial tension to almost zero to disperse multiple droplets even in the thermodynamic equilibrium (microemulsion) (25,36). The surfactant monolayer acts as a kinetic barrier to suppress the coalescence and decreases the hydrostatic pressure of the droplet to suppress the coarsening (9,25). These features are analogous to the terminal regions of pre-rRNAs at the surfaces of FCs. The fact that some RNAs and proteins can act as surfactants was discussed in recent researches (37,38). There are also differences between nascent pre-rRNA and surfactants. First and foremost, surfactants are localized at interfaces at the thermodynamic equilibrium, while the terminal regions of nascent pre-rRNAs are localized at the FC-DFC interfaces only during the transcription and their surface activity is regulated by Pol I transcription dynamics. Second, the hydrophobic chains of surfactants in a liquid monolayer tend to form a melt, while the terminal regions of pre-rRNA are extended due to the RBPs freely diffusing in the DFC layers. Last, but not least, surfactants can freely diffuse in a monolayer, while the terminal regions of nascent pre-rRNAs are anchored to the rDNA at the interface via Pol I and thus are not mobile.

In our model, we focused on FBLs because these RBPs contribute most significantly to the localization of the terminal regions of pre-rRNAs to the DFC layer (13). FRAP experiments on the condensates of FBLs reconstituted *in vitro* suggest that these condensates are thixotropic (and probably also be viscoelastic) and become gels in a long timescale (10). Indeed, recent experiments showed that long non-coding RNA SLERT facilitates the transition of the RNA helicase DDX21 to the closed conformation and ensures the fluidity of proteins in the DFC layer (39). This mechanism ensures the validity of our assumption that unbound FBLs can diffuse freely in the DFC layer, although we did not take into account these factors explicitly.

Although our theory probably could capture the essential features that are necessary to understand the assembly mechanism of the multiphase structure of nucleolus, it is not complete. First, the layers at the interface between FCs and GC are not uniform, but DFC regions are flanked by the regions, in which FBLs are not observed by super-resolution microscopy and are not visible by electron microscopy. Second, we did not take into account the downstream pre-rRNA regions that are not bound by FBLs. These downstream regions were observed at the surfaces of FCs (13). The direct contribution of rDNAs localized at the surfaces of FCs are not significant, see S4 in the SI Appendix. Third, we did not explicitly take into account the transport dynamics of pre-rRNAs in GCs. Fourth, RNAs can be not only cleaved, but also can be folded and/or chemically modified during the co-transcriptional processing. The processing of pre-rRNAs in yeast has been well-documented (40,41), but much less in higher eukaryotes (42,43), in which the multiphase structure of nucleolus is usually observed. It is of interest to understand the mechanism of the processed pre-rRNAs, which have the affinity to FCs, penetrate through the DFC layer towards the GC. One possible hypothesis is that the processed pre-rRNAs pass the flanking regions of the DFC layer. These flanking regions increase the density of the terminal regions of pre-rRNAs in the DFC compartments and enhance the lateral osmotic pressure. The enhanced lateral pressure and/or the kinetic trapping of the coalescence of FCs by nascent pre-rRNAs may explain the fact that the radius of FCs observed in experiments is smaller than that predicted by our theory. Whether the multiphase structure of the nucleolus contributes to its function is not yet clear. It is of interest to extend our theory to understand the structure-function relationship of the nucleolus (44). However, following the approach of the classical theoretical physics, our present theory should be experimentally tested before adding more assumptions that have not been experimentally explored.

The key assumption of our theory is that the terminal regions of nascent pre-rRNAs span between the top to the bottom of the DFC layer. This is motivated by a recent experiment that the upstream half of ETSs are localized at the DFC layer and the downstream half is localized at the FC (13). The stretching of the terminal regions results from the multivalent interactions between FBLs and thus our theory should be tested in the condition of high FBL volume fraction. In the conventional picture, the DFC layer is composed of 5’ ETSs that are already cleaved and folded by RNA processing. Such diffusive molecules tend to make the interaction between condensates attractive (the depletion interaction) (25), however, the non-equilibrium nature of the transcription may change the situation. We envisage a future theoretical study to predict the stable size of FCs with the conventional picture to check if the latter agrees with existing experiments, including ours.

Our theory may provide insight into the size control mechanism of other nuclear bodies. The size of nuclear speckles (5) increases by the suppression of transcription. The stability of transcriptional condensate depends on the frequency of transcription and the length of transcripts (45,46). Transcriptional condensates are actively studied in recent experiments (45-49). The simulations by Henninger and coworkers predict that RNAs diffuse freely in the interior of condensates and destabilize the condensates due to the electrostatic repulsion (45). In contrast, Hilbert and coworkers argued that DNA-Pol II-RNA complexes are localized at the interface between transcriptional condensates (rich in RNA and proteins) and the exterior (rich in DNA) and act as surfactants that suppress the coalescence and coarsening of the condensates (46), analogous to microemulsion (see above). Our theory is somewhat similar to the idea of Hilbert and coworkers, but different in the fact that the complexes of nascent pre-rRNAs and RBPs form distinct phase at the interface between FCs and GC due to the transcription and FCs and GC are somewhat similar in terms of the relative affinity to the nucleoplasm, as characterized by the interfacial tension (10). The Pol II transcription is regulated by the phosphorylation of the C-terminal domains and is somewhat different from the Pol I transcription. It is of interest to take into account such differences in an extension of our present theory to understand the assembly mechanism of nuclear bodies and their functions.

The multiphase structure of nucleoli may be somewhat analogous to that observed in nuclear stress bodies (nSBs) by electron microscopy (50). nSBs are assembled during the thermal stress condition and are composed of HSATIII architectural RNA and specific RBPs (51). HSATIII is transcribed by Pol II from the pericentromeric Satellite III regions, which are enriched in tandem repeats (52,53). These features common with nucleoli motivate us to think of a general mechanism involved in the assembly of the multiple subcompartments. The nascent RNAs, which are produced by transcription and are modulated by RNA processing, are important elements that regulate the multiphase structures, and possibly their functions, of nuclear bodies.

## MATERIALS AND METHODS

### Transcription and RNA processing

The occupancy *n*_ps_ of the TSS of the active rDNA by Pol I follows the kinetic equation

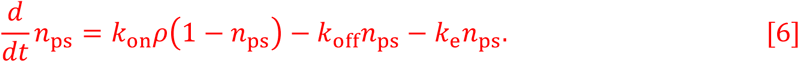

Eq. 6 suggests that the occupancy *n*_ps_ changes due to the binding of Pol I from the FC (the first term), the unbinding of Pol I from the TSS to the FC without starting transcription (the second term), and the initiation of the elongation of Pol I (the third term), see Fig. 2A. *k*_on_ is the rate constant for the binding of Pol I to TSS and *ρ* is the concentration of Pol I in a FC (which will be determined later). *k*_off_ is the rate constant for the unbinding of Pol I from TSS. *k*_e_ is the rate of the transition of Pol I from the bound state to the elongation state.

The number *n*_pe_ of elongating Pol I protein complexes with pre-processed nascent pre-rRNA follows the kinetic equation

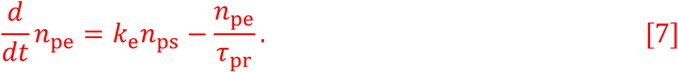

Eq. 7 suggests that the number *n*_pe_ changes due to the release of Pol I from TSS for elongation (the first term) and the cleavage of the FBL-binding terminal region of nascent pre-rRNA (the second term), see Fig. 2B. *τ*_pr_ is the average time between the transcription initiation and the cleavage of the terminal regions of the pre-rRNA.

The number *n*_p0_ of elongating Pol I with processed nascent pre-rRNA follows the kinetic equation

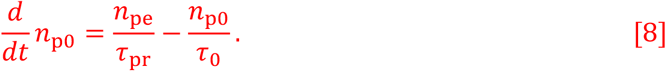

Eq. 8 suggest that the number *n*_p0_ changes due to the the cleavage of the terminal regions of pre-rRNAs (the first term) and the transcription termination (the second term), see Fig. 2C. *τ*_0_ is the average time between the cleavage of FBL-binding region and the transcription termination. The numbers, *n*_ps_, *n*_pe_, and *n*_p0_, of Pol I in each state are derived for the steady state, 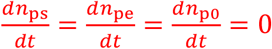, by using eqs. 6-8. The inverse time *k* of the transcription initiation is much smaller than the inverse *k*_off_ of the unbinding of Pol I from TSS, *k*_e_ ≪*k*_off_. The concentration *ρ* of Pol I in a FC is determined by the condition

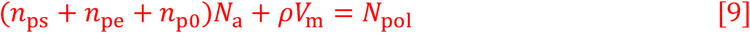

that the sum of the number of Pol I during transcription (the first term) and the number of Pol I diffusing in microphases (the second term) in the system is constant. All the parameters involved in eq. 9 do not depend on the size of FCs, see eqs. 6-8. The concentration *ρ* of Pol I in FCs thus does not depend on the size of FCs.

The surface density *σ*_in_ of the FBL-binding terminal regions of nascent pre-rRNAs at each FC (the number of nascent pre-rRNAs per unit area of each FC surface) is represented as

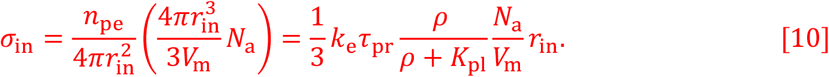

by assuming that the terminal regions of nascent pre-rRNA is distributed uniformly at the surfaces of FCs. We note that 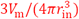 is the number of FCs in a nucleolus. The last form of eq. 10 is derived by using the solution of eqs. 6-8.

### Thickness of DFC layer

The exterior radius *r*_ex_ (see also Fig. 1B) is determined by the relationship

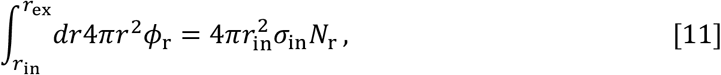

where *N*_r_ is the average number of units in the terminal region of a nascent pre-rRNA. It is a mean field treatment that assumes that nascent RNAs are composed of the same number of units and is effective within the Alexander-de Gennes approximation, with which the brush height is determined by the distance between neighboring grafting points and the average number of units per chain (if one neglects the fact that the lateral fluctuations of a chain composed of *N*_r_ units is limited to 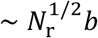) (21, 54).

### Free energy of DFC layer

The free energy of each DFC layer has the form

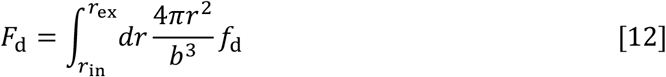

With

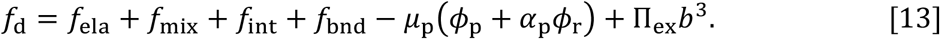

Eq. 12 is the functional of the local volume fraction *ϕ*_r_ of nascent pre-rRNA units, the local volume fraction *ϕ*_p_ of RBPs, and the occupancy *α*_p_ of nascent pre-rRNA units by RBPs, which are functions of the distance *r* from the center of the FC (*r*_in_ < *r*< *r*_ex_), see Fig. 1B. *b* is the length of a pre-rRNA unit. This free energy is composed of 4 contributions: *f*_ela_ is the elastic free energy density of the terminal regions of nascent RNAs, *f*_mix_ is the free energy density due to the mixing entropy of RBPs and solvent molecules, *f*_int_ is the free energy density due to the interactions between RBPs, and *f*_bnd_ is the free energy density due to the binding of RBPs to the terminal regions of nascent RNAs. *µ*_p_ is the chemical potential of RBPs and ∏_ex_ is the osmotic pressure from the exterior.

The (entropic) elastic free energy density has the form

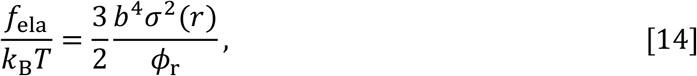

where *σ*(*r*) is the surface density of FBL-binding regions of nascent pre-rRNAs. 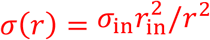 for cases in which the surface of FCs is sphere and *σ*(*r*) = *σ*_in_ for cases in which the surface of FCs is planer (which corresponds to the limit of 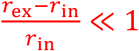). The volume fraction *ϕ*_r_ of pre-rRNA units decreases as the terminal regions of the nascent pre-rRNAs are stretched. Eq. 14 represents the fact that the elastic free energy *f*_ela_ increases as these terminal regions are stretched. Eq. 14 takes into account the spherical geometry of the system in an extension of the Alexander model (that assumes that the concentration of the terminal regions of nascent pre-rRNA is uniform in the DFC layer in the limit of 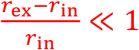) in the spirit of the Daoud-Cotton theory (55), see sec. S5 in the SI Appendix for the derivation. *k*_B_ is the Boltzmann constant and *T* is the absolute temperature.

The free energy due to the mixing entropy of RBPs and solvent molecules has the form

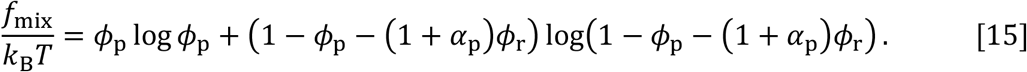

The mixing free energy, eq. 15, has the contributions of the mixing entropy of RBPs (the first term) and the mixing entropy of solvent molecules (the second term).

The free energy due to the interactions between RBPs has the form

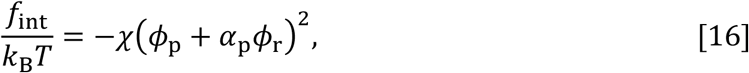

where *χ* is the interaction parameter that accounts for the attractive interactions. For simplicity, we assumed that RBPs bound to the terminal regions of nascent pre-rRNAs are equivalent to RBPs freely diffusing in the DFC layer and that solvent molecules are equivalent to nascent RNA units in terms of the interactions, see also the discussion in Parameter estimate.

The free energy due to the binding of RBPs to nascent RNAs has the form

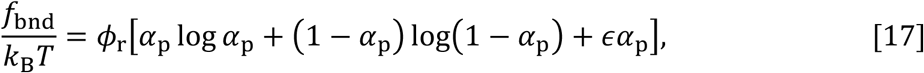

where *εk*_B_*T* is the energy increase due to the binding of RBPs to nascent RNAs. For simplicity, we assumed that each nascent RNA unit has one binding site of RBPs.

### Free energy of system

The free energy *F* of the system, eq. 2 is derived by using the fact that the number of microphases in the system is 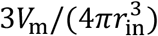 to derive this form. The interfacial tension between GC and nucleosol is relative small (3,10) and FC is excluded out from the nucleolus by the transcription inhibition (3,11), implying that the interfacial tension between FC and nucleosol is even smaller. In vitro experiments suggest that the attractive interaction between FBLs is much larger than that between nucleophosmins, a major component of GC (10). It implies that the interfacial tension between DFC and GC and that between DFC and FC are larger than their difference. By neglecting the difference between these interfacial tensions, the interfacial tensions *γ*_in_ and *γ*_ex_ at the interior and the exterior interfaces are represented by the forms

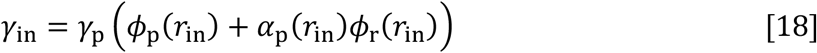

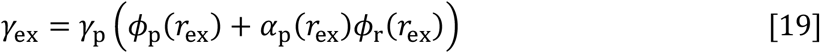

where *γ*_p_ is the surface tension between a nucleosol and a liquid of the RBPs and is proportional to *χ*.

### Lateral osmotic pressure

The lateral osmotic pressure ∏_∥_ can be defined by

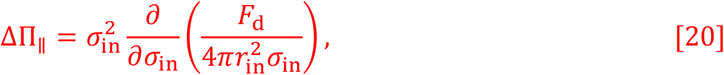

where the derivative with respect to *σ*_0_ is taken with the condition that 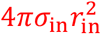 is constant (this increases the area of the surface of a FC without changing the composition of the DFC layer). Δ∏_∥_ is the lateral osmotic pressure, from which the contribution of the isotropic osmotic pressure is subtracted.

### Radius of FCs at free energy minimum

The free energy, eq. 1, is represented as a function only of the radius *r*_in_ of FCs if the occupancy, *α*_p_, and the volume fractions, *ϕ*_p_ and *ϕ*_r_, derived by using eqs. 2-5 are substituted into eq. 1. The radius *r*_in_ at the steady sate is derived by the condition 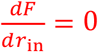 with the condition that *σ*_0_ ∝ *r*_in_ (this increases the area of the surfaces of FCs with the optimization of the arrangement of the active rDNA repeats). This leads to the form

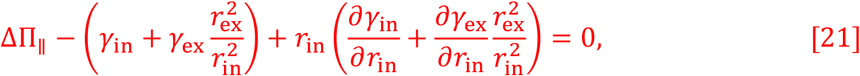

which represent the fact that the surface tension is balanced to the surface pressure generated by the FBRs of nascent pre-rRNAs.

### Cell culture, drug treatment, reverse transcription-quantitative PCR (RT-qPCR), immunofluorescence, and quantification of the size of FCs

HeLa cells were maintained in DMEM containing high glucose (Nacalai Tesque, Cat# 08458-16) supplemented with 10% FBS (Sigma) and Penicillin-Streptomycin (Nacalai Tesque). Cells were treated with BMH-21 (Selleckchem, Cat# S7718) or CX-5461 (AdooQ Bioscience) for 3 hours. Quantification of nascent pre-rRNAs (forward primer: 5’-CCTTCCCCAGGCGTCCCTCG-3’, reverse primer: 5’-

GGCAGCGCTACCATAACGGA-3’) (11) by RT-qPCR was performed using LightCycler 480 II (Roche). GAPDH mRNAs were used as a loading control (14). The antibodies to UBF (Santacruz, F-9, sc-13125, mouse monoclonal antibody, 1:50 dilution), FBL (Proteintech, 16021-1-AP, rabbit polyclonal antibody, 1:500 dilution), and NPM1/Nucleophosmin (Abcam, ab183340, SP236, rabbit polyclonal antibody, 1:100 dilution) were used to visualize FCs, DFCs, and GCs in immunofluorescence, respectively. Cells were grown on coverslips (Matsunami; 18 mm round) and fixed with 4% paraformaldehyde/PBS at room temperature for 10 min. Then, the cells were washed three times with 1 x PBS, permeabilized with 0.5% Triton-X100/PBS at room temperature for 5 min, and washed three times with 1 x PBS. The cells were incubated with 1 x blocking solution (Roche, Blocking reagent and TBST [1 x TBS containing 0.1% Tween 20]) at room temperature for 1 hour. Then, the coverslips were incubated with primary antibodies in 1 x blocking solution at room temperature for 1 hour, washed three times with TBST for 5 min, incubated with secondary antibodies (anti-mouse IgG, Alexa Fluor 488, superclonal [Thermo Fisher Scientific, Cat#A28175], anti-rabbit IgG, Alexa Fluor 568 [Thermo Fisher Scientific, Cat#A11036]) at room temperature for 1 hour, and washed three times with TBST for 5 min. The coverslips were mounted with a Vectashiled hard set mounting medium with DAPI (Vector, H-1500). Super-resolution images were acquired using ZEISS LSM900 with Airyscan 2. Quantification of the FCs marked by UBF staining within the nucleoli labeled by NPM1 staining was performed using NIS Elements Advanced Research (NIKON). “FillArea” and “MaxFeret” of the UBF foci (FCs) detected with an intensity threshold were quantified as area and Lx of the FCs (Fig. S4).

## DATA AVAILABILITY

Mathematica file used for the numerical calculations are available in figshare with the identier (https://doi.org/10.6084/m9.figshare.16599446).

## ACKNOWLEDGMENT

This research was supported by KAKENHI grants from the Ministry of Education, Culture, Sports, Science, and Technology (MEXT) of Japan [to T. Yamamoto (20H05934, 21K03479, 21H00241), T. Yamazaki (21H00253, 22H02545), KN (19K06478, 22K06083),and TH (20H00448, 20H05377,21H05276)], JST, PRESTO Grant Number JPMJPR18KA (to T. Yamamoto), the Mochida Memorial Foundation for Medical and Pharmaceutical Research (to T. Yamazaki), the Naito Foundation (to T. Yamazaki), the Takeda Science Foundation (to T. Yamazaki), and JST CREST Grant Number JPMJCR20E6 (to T.H.). T. Yamazaki thanks Toyofumi Kameoka (NIKON SOLUTIONS CO., LTD.) and Takayuki Funato (NIKON SOLUTIONS CO., LTD.) for support of microscopic image analyses. T. Yamamoto acknowledges the fruitful discussion with Noriko Saito (Japanese Foundation for Cancer Research), Yuma Ito (Tokyo Institute of Technology), Satoru Ide (National Institute of Genetics), Tsutomu Suzuki (Univ. of Tokyo), Takahiko Kobayashi (Univ. of Tokyo), Yutetsu Kuruma (JAMSTEC), Shintaro Iwasaki (Univ. of Tokyo), Sumio Sugano (Chiba University), Haruhiko Siomi (Keio Univ.), Hideaki Matsubayashi (Johns Hopkins Univ.), and Naomichi Takemata (Kyoto Univ.).

## FIGURE LEGENDS

**Figure 1 Multiphase structure of a nucleolus.**(A) A nucleolus is composed of multiple fibrillar center (FC) microphases in the sea of the granular component (GC). There is a layer of dense fibrillar component (DFC) between each FC and GC. (B) RNA polymerase I (Pol I) molecules (white particles) are entrapped in FC microphases (light blue) and the active rDNA units (black line) are localized at the surfaces of microphases. Nascent pre-rRNAs (green particles) are thus at the surfaces of the microphase and form a DFC layer with RNA-binding proteins (magenta particles). The interface between FC and DFC is located at a distance *r*_in_ from the center and the interface between DFC and GC is located at a distance *r*_ex_.

**Figure 2 Model of transcription dynamics.** DNA (black solid line) is localized at the surface of an FC (cyan). (A) RNA polymerase I (Pol I) in a microphase binds to the transcription starting site (TSS) of an active rDNA repeat unit. The bound Pol I starts transcription with the rate *k*_e_ or returns to the microphase without starting transcription. During the transcription, Pol I migrates uni-directionally towards the transcription terminating site (TTS) while polymerizing a nascent pre-rRNA. The terminal region of the nascent pre-ribosomal RNA (pre-rRNA), to which fibrillarin binds, are cleaved by the co-transcriptional RNA processing with the rate 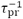. The terminal region is released to GC. After the cleavage of the terminal region, Pol I continues transcription until it reaches the TTS. At the TTS, Pol I is releaed to the FC with the rate 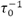.

**Figure 3 Composition of planer DFC layer vs interaction parameter *χ*.** (A) The volume fraction *ϕ*_r_ of pre-rRNA units (green), the volume fraction *ϕ*_p_ of freely diffusing RBPs (magenta), the volume fraction *ϕ*_s_ of solvent molecules (cyan) in a DFC layer are shown as functions of the interaction parameter *χ*. (B) The occupancy *α*_p_ of pre-rRNA terminal region by RBPs is shown as a function of the interaction parameter *χ*. We used *µ*_*p*_/(*k*_B_*T*) = −10.0, *ε* = −12.0, *σ*_in_*b*^2^ = 0.05, and ∏_ex_*b*^3^/(*k*_B_*T*) = 0.0 for the calculations, see also Table 2. The solid lines are derived by numerically solving eqs. S15, S16, and S20 in the SI Appendix. The broken lines are derived by using eqs. S47 (shown for *χ* > 10.1).

**Figure 4 Profile of volume fraction *ϕ*_r_ of pre-rRNA units in DFC layer.** The volume fraction of nascent pre-rRNA units is shown as a function of the position *r* in the DFC layer (*r*_in_ < *r*< *r*_ex_). The solid dark green line is derived by numerically calculating eqs. S13, S15, and S16 in the SI Appendix for *ζ* = 0.06 with the condition that the volume fraction of solvent is zero and the occupancy of the terminal regions of pre-rRNAs by RBPs is unity. The values of other parameters are summarized in Table 2. The broken light green line is derived by using eq. S59 in the SI Appendix with *ϕ*_ex_ = 0.0157 and *r*_ex_ = 1.98 (which were derived from the numerical calculation to obtain the light green line). The volume fraction of freely diffusing RBPs is shown in the inset.

**Figure 5 Free energy *F* of nucleolus.** The free energy *F* of the system is shown as a function of the radius *r*_in_ of FCs (A) and the ratio *r*_ex_/*r*_in_ of the external radius to the internal radius (B). The black solid line is the total free energy, including the free energy of DFC layers (shown by the magenta broken line, the first term of eq. 2) and the surface free energy (shown by the cyan broken line, the second and third terms of eq. 2) for *ζ* = 0.06. The values of other parameters are summarized in Table 2.

**Figure 6 Radius *r*_ii_ of FCs vs rescaled transcription rate *ζ*.** The radius *r*_in_ of FCs at the free energy minimum is shown as a function of rescaled transcription rate *ζ*. The solid dark green line is derived by numerically calculating eqs. S13, S15, and S16 in the SI Appendix with the condition that the volume fraction of solvent is zero and the occupancy of the terminal regions of pre-rRNAs by RBPs is unity. The orange broken line is derived by using eq. 5. The parameters used for the calculations are summarized in Table 2.

**Figure 7 Mild Pol I inhibition by BMH-21 increases the size of FCs.** (A) Immunofluorescence of UBF (FC) and NPM1 (GC) in HeLa cells with or without BMH-21 treatments. Scale bar, 10 µm. (B and C) Quantification of longest axis (B) and area (C) of the FCs in cells under indicated conditions. Each scatter dot plot shows the mean (black line). Dots indicate all points of quantified data (n = 250). Mean longest axes of the FCs is shown below: 0 µM: 0.4907 µm, 0.0625 µM: 0.7440 µm, 0.125 µM: 0.8710 µm, 0.25 µM: 1.131 µm. Mean areas of the FCs are shown below: 0 µM: 0.170 µm^2^, 0.0625 µM: 0.3255 µm^2^, 0.125 µM: 0.4347 µm^2^, 0.25 µM: 0.7312 µm^2^. Statistical analyses using Kruskal-Wallis test with Dunn’s multiple comparison test were performed and the results are shown as follows. (B) 0 µM vs 0.0625 µM: P < 0.0001, 0 µM vs 0.125 µM: P < 0.0001, 0 µM vs 0.25 µM: P < 0.0001, 0.0625 µM vs 0.125 µM: P < 0.0001, 0.0625 µM vs 0.25 µM: P < 0.0001, 0.125 µM vs 0.25 µM: P = 0.0008. (C) 0 µM vs 0.0625 µM: P = 0.0003, 0 µM vs 0.125 µM: P < 0.0001, 0 µM vs 0.25 µM: P < 0.0001, 0.0625 µM vs 0.125 µM: P < 0.0001, 0.0625 µM vs 0.25 µM: P < 0.0001, 0.125 µM vs 0.25 µM: P = 0.0010. (D and E) Graphs showing the mean longest axis (D) and area (E) of the FCs with SEM vs pre-rRNA expression levels. The pre-rRNA expression level in untreated cells is defined as 1.

**Figure 8 Exponent that accounts for the dependence of the size of FCs on pre-rRNA expression level.** The data in Fig. 7E was shown in the double-logarithm plot and fitted with a power function. The magenta and cyan dots are the results by suppressing the Pol I transcription by using BMH-21 and CX-5461. The slope of the double-logarithm plot is the exponent that accounts for the dependence of the radius of FCs on the transcription level. The curve fitting shows that the exponent is -0.49 for the case of the BMH-21 treatment, -0.46 for the case of the CX-5461, and -0.48 if both data are fitted.

**Figure 9 Summary of results.** RNP complexes enhance or suppress the growth of condensates depending on whether the RNP complexes are mobile in the interior or tethered to the surfaces of the condensates. A. The multivalent interaction between the RNP complexes enhance the growth of the condensates if these condensates are assembled by the RNP complexes. B. The multivalent interaction between the RNP complexes suppress the growth of the condensates if these complexes are tethered to the surfaces of the condensates assembled by other RNAs and proteins.

## LEGENDS OF SUPPLEMENTARY FIGURES

**Figure S1 BMH-21 and CX-5461 treatment reduce pre-rRNA expression levels in a dose-dependent manner**.

(*A*) Quantification of pre-rRNA expression levels by RT-qPCR in BMH-21-untreated and -treated conditions. Data are represented as mean ± SD (*n* =3). (*B*) Quantification of pre-rRNA expression levels by RT-qPCR in CX-5461-untreated and -treated conditions. Data are represented as mean ± SD (*n* =3).

**Figure S2 FCs and DFCs in the BMH-21 and CX-5461-untreated and -treated cells**. (*A* and *B*) Immunofluorescence of UBF (FC) and FBL (DFC) in HeLa cells with or without BMH-21 (*A*) or CX-5461 (*B*) treatments. Scale bar, 10 µm.

**Figure S3 Mild Pol I inhibition by CX-5461 increases the size of FCs**.

(*A*) Immunofluorescence of UBF (FC) and NPM1 (GC) in HeLa cells with or without CX-5461 treatments. In the cells treated with 2 µM CX-5461, the cells with large granules (upper) or nucleolar caps (lower) were observed. Scale bar, 10 µm. (*B* and *C*) Quantification of longest axis (*B*) and area (*C*) of the FCs in cells under indicated conditions. Each scatter dot plot shows the mean (black line). Dots indicate all points of quantified data (*n* = 500). Mean longest axes of the FCs is shown below: 0 µM: 0.5196 µm, 0.25 µM: 0.6367 µm, 0.5 µM: 0.7192 µm, 1 µM: 1.059 µm. Mean areas of the FCs are shown below: 0 µM: 0.1155 µm^2^, 0. 25 µM: 0.2171 µm^2^, 0.5 µM: 0.3058 µm^2^, 1 µM: 0.6544 µm^2^. Statistical analyses using Kruskal-Wallis test with Dunn’s multiple comparison test were performed and the results are shown as follows. (*B*) 0 µM vs 0.25 µM: *P* < 0.0001, 0 µM vs 0.5 µM: *P* < 0.0001, 0 µM vs 1 µM: *P* < 0.0001, 0.25 µM vs 0.5 µM: *P* = 0.0211, 0.25 µM vs 1 µM: *P* < 0.0001, 0.5 µM vs 1 µM: *P* < 0.0001. (*C*) 0 µM vs 0.25 µM: *P* < 0.0001, 0 µM vs 0.5 µM: *P* < 0.0001, 0 µM vs 1 µM: *P* < 0.0001, 0.25 µM vs 0.5 µM: *P* = 0.0012, 0.25 µM vs 1 µM: *P* < 0.0001, 0.5 µM vs 1 µM: *P* < 0.0001. (*D* and *E*) Graphs showing the mean longest axis (*D*) and area (*E*) of the FCs with SEM vs pre-rRNA expression levels. The pre-rRNA expression level in untreated cells is defined as 1.

**Figure S4 Quantification of the FCs in the nucleoli**

Screenshots of detections of FCs and GCs by NIS Elements Advanced Research software (NIKON) are shown. Green circles indicate UBF foci, and red circles indicate nucleoli labeled by NPM1 antibodies. UBF foci detected in the nucleoli (lower right) were quantified.

