## Supplemental File for "Lateral surface pressure generated by nascent ribosomal RNA suppresses growth of fibrillar centers in the nucleolus"

**This PDF file includes:**

Supplementary text
Table S1 (not allowed for Brief Reports)
SI References

### Supporting Information Text

#### S1. Minimum of free energy

**A. Free energy.** The free energy of the system has the form

$$\frac{F}{V_{\rm m}} = \frac{3}{4\pi r_{\rm in}^3}\left[F_{\rm d} + 4\pi r_{\rm in}^2\gamma_{\rm in} + 4\pi r_{\rm ex}^2\gamma_{\rm ex}\right]. \quad \text{[S1]}$$

$F_{\rm d}$ is the free energy of a DFC layer and is a functional of the occupancy $\alpha_{\rm p}$ of the fibrillarin-binding region (FBR) of nascent pre-rRNA, the volume fraction $\phi_{\rm p}$ of fibrillarin (FBL) RNA-binding proteins (RBPs), and the volume fraction $\phi_{\rm r}$ of the nascent pre-rRNA units (see below). $\gamma_{\rm in}$ is the interfacial tensions at the FC-DFC interface and $\gamma_{\rm ex}$ is the interfacial tension at the DFC-GC interface. $r_{\rm in}$ is the distance between the center of FC and the FC-DFC interface. $r_{\rm ex}$ is the distance between the center of FC and the DFC-GC interface.

The free energy of a DFC layer is written in the form

$$F_{\rm d} = \int_{r_{\rm in}}^{r_{\rm ex}} \frac{4\pi r^2 dr}{b^3} f_{\rm d}, \quad \text{[S2]}$$

where $f_{\rm d}$ is the free energy density in the DFC layer and $r$ is the radical coordinate from the center of the FC. $b$ is the length of a pre-rRNA unit. The free energy density $f_{\rm d}$ is composed of 4 contributions

$$f_{\rm d} = f_{\rm ela} + f_{\rm mix} + f_{\rm int} + f_{\rm bnd} - \mu_{\rm p}(\phi_{\rm p} + \alpha_{\rm p}\phi_{\rm r}) + \Pi_{\rm ex} b^3, \quad \text{[S3]}$$

where $f_{\rm ela}$ is the elastic free energy density of nascent RNA transcripts, $f_{\rm mix}$ is the free energy density due to the mixing entropy of RBPs and solvent molecules, $f_{\rm int}$ is the free energy density due to the interactions between RBPs, and $f_{\rm bnd}$ is the free energy density due to the binding of RBPs to nascent RNA transcripts. $\mu_{\rm p}$ is the chemical potential of RBPs and $\Pi_{\rm ex}$ is the external pressure. The external radius $r_{\rm ex}$ and the volume fraction $\phi_{\rm r}$ of nascent pre-rRNA have the relationship

$$\int_{r_{\rm in}}^{r_{\rm ex}} dr\, 4\pi r^2 \phi_{\rm r} = 4\pi r_{\rm in}^2 \sigma_{\rm in} N_{\rm r}. \quad \text{[S4]}$$

The elastic free energy density has the form

$$\frac{f_{\rm ela}}{k_{\rm B}T} = \frac{3}{2}\frac{b^4\sigma^2(r)}{\phi_{\rm r}}, \quad \text{[S5]}$$

where $\sigma(r) = \sigma_{\rm in} r_{\rm in}^2/r^2$ for the spherical geometry and $\sigma(r) = \sigma_{\rm in}$ for the planer geometry. $k_{\rm B}$ is the Boltzmann constant and $T$ is the absolute temperature. The derivation of eq. S5 is shown in sec. S5 in this SI Appendix.

The free energy due to the mixing entropy of RBPs and solvent molecules has the form

$$\frac{f_{\rm mix}}{k_{\rm B}T} = \phi_{\rm p}\log\phi_{\rm p} + (1-\phi_{\rm p}-(1+\alpha_{\rm p})\phi_{\rm r})\log(1-\phi_{\rm p}-(1+\alpha_{\rm p})\phi_{\rm r}). \quad \text{[S6]}$$

The free energy due to the interactions between RBPs has the form

$$\frac{f_{\rm int}}{k_{\rm B}T} = -\chi(\phi_{\rm p}+\alpha_{\rm p}\phi_{\rm r})^2, \quad \text{[S7]}$$

where $\chi$ is the interaction parameter that accounts for the attractive interaction between RBPs. For simplicity, we assumed that RBPs that are bound to nascent RNA transcripts are equivalent to RBPs that are freely diffusing in the DFC layer. The solvent molecules (water) have affinity to nascent RNA units rather than RPBs. We thus assume that solvent molecules and nascent pre-rRNA units are equal in terms of the interaction.

The free energy due to the binding of RBPs to nascent RNA transcripts has the form

$$\frac{f_{\rm bnd}}{k_{\rm B}T} = \phi_{\rm r}\left[\alpha_{\rm p}\log\alpha_{\rm p} + (1-\alpha_{\rm p})\log(1-\alpha_{\rm p}) + \epsilon\alpha_{\rm p}\right], \quad \text{[S8]}$$

where $-\epsilon k_{\rm B}T$ is the energy increase due to the binding of RBPs to nascent RNA transcripts. For simplicity, we assumed that each nascent RNA unit has one binding site of RBPs.

**B. Local equilibrium.** With the variations $\alpha_\mathrm{p}(r) \to \alpha_\mathrm{p}(r) + \delta\alpha_\mathrm{p}(r)$, $\phi_\mathrm{p}(r) \to \phi_\mathrm{p}(r) + \delta\phi_\mathrm{p}(r)$, and $\phi_\mathrm{r}(r) \to \phi_\mathrm{r}(r) + \delta\phi_\mathrm{r}(r)$, the free energy changes as

$$\frac{\delta F}{V_\mathrm{m}} = \frac{3}{4\pi r_\mathrm{in}^3}\int_{r_\mathrm{in}}^{r_\mathrm{ex}} dr\, \frac{4\pi r^2}{b^3}\left[\left(\frac{\delta f_\mathrm{d}}{\delta\alpha_\mathrm{p}(r)}\delta\alpha_\mathrm{p}(r) + \frac{\delta f_\mathrm{d}}{\delta\phi_\mathrm{p}(r)}\delta\phi_\mathrm{p}(r) + \frac{\delta f_\mathrm{d}}{\delta\phi_\mathrm{r}(r)}\delta\phi_\mathrm{r}(r)\right)\right] + \delta r_\mathrm{ex}\frac{\partial}{\partial r_\mathrm{ex}}\left(\frac{F}{V_\mathrm{m}}\right). \quad \text{[S9]}$$

Because the free energy density $f_\mathrm{d}$ does not include spatial derivatives of $\alpha_\mathrm{p}(r)$, $\phi_\mathrm{p}(r)$, and $\phi_\mathrm{r}(r)$, the functional derivatives of $f_\mathrm{d}$ with respect to $\alpha_\mathrm{pl}(r)$, $\phi_\mathrm{pl}(r)$, and $\phi_\mathrm{r}(r)$ can be replaced to the partial derivatives of $f_\mathrm{d}$ with respect to $\alpha_\mathrm{p}$, $\phi_\mathrm{p}$, and $\phi_\mathrm{r}$. The variation $\delta r_\mathrm{ex}$ of the external radius results from the variation $\delta\phi_\mathrm{r}(r)$ with the condition of eq. S4:

$$\delta r_\mathrm{ex} = -\frac{b^3}{4\pi r_\mathrm{ex}^2}\frac{1}{\phi_\mathrm{r}(r_\mathrm{ex})}\int_{r_\mathrm{in}}^{r_\mathrm{ex}} dr\, \frac{4\pi r^2}{b^3}\delta\phi_\mathrm{r}(r). \quad \text{[S10]}$$

At the minimum of the free energy (the local equilibrium), $\delta F = 0$ for any functions of $\delta\alpha_\mathrm{pl}(r)$, $\delta\phi_\mathrm{pl}(r)$, and $\delta\phi_\mathrm{r}(r)$:

$$\frac{\partial f_\mathrm{d}}{\partial \alpha_\mathrm{p}} = 0 \quad \text{[S11]}$$

$$\frac{\partial f_\mathrm{d}}{\partial \phi_\mathrm{p}} = 0 \quad \text{[S12]}$$

$$\frac{\partial f_\mathrm{d}}{\partial \phi_\mathrm{r}} = \mu_\mathrm{r} \quad \text{[S13]}$$

with

$$\mu_\mathrm{r} = \left.\frac{f_\mathrm{d}}{\phi_\mathrm{r}}\right|_{r\to r_\mathrm{ex}} + \left(\frac{2b^3\gamma_\mathrm{ex}}{r_\mathrm{ex}} + \Pi_\mathrm{ex}b^3\right)\frac{1}{\phi_\mathrm{r}(r_\mathrm{ex})}. \quad \text{[S14]}$$

By using eqs. S3 - S8, eq. S12 is rewritten in the form

$$\frac{\mu_\mathrm{p}}{k_\mathrm{B}T} = \log\phi_\mathrm{p} - \log(1-\phi_\mathrm{p}-(1+\alpha_\mathrm{p})\phi_\mathrm{r}) - 2\chi(\phi_\mathrm{p}+\alpha_\mathrm{p}\phi_\mathrm{r}). \quad \text{[S15]}$$

Eq. S15 suggests that the chemical potential of RBPs is uniform in the DFC layer. By using eqs. S3 - S8, eq. S11 is rewritten in the form

$$\log\alpha_\mathrm{p} - \log(1-\alpha_\mathrm{p}) + \epsilon - 1 - \log\phi_\mathrm{p} = 0, \quad \text{[S16]}$$

We eliminated $\mu_\mathrm{p}$ in eq. S11 by using eq. S15 to derive the form of eq. S16. By using eqs. S3 - S8, eq. S13 is rewritten in the form

$$\frac{\mu_\mathrm{r}}{k_\mathrm{B}T} = -\frac{3}{2}\frac{(\sigma^2(r)b^2)^2}{\phi_\mathrm{r}^2} - (1+\alpha_\mathrm{p})\log(1-\phi_\mathrm{p}-(1+\alpha_\mathrm{p})\phi_\mathrm{r}) - (1+\alpha_\mathrm{p}) - 2\chi\alpha_\mathrm{p}(\phi_\mathrm{p}+\alpha_\mathrm{p}\phi_\mathrm{r})$$
$$+\alpha_\mathrm{p}\log\alpha_\mathrm{p} + (1-\alpha_\mathrm{p})\log(1-\alpha_\mathrm{p}) + \epsilon\alpha_\mathrm{p} - \frac{\mu_\mathrm{p}}{k_\mathrm{B}T}\alpha_\mathrm{p}. \quad \text{[S17]}$$

Eq. S13 is integrated with respect to $\phi_\mathrm{r}$ as

$$f_\mathrm{d} - \mu_\mathrm{r}\phi_\mathrm{r} = -\Pi b^3, \quad \text{[S18]}$$

where $-\Pi b^3$ is the integral constant. By eliminating $\mu_\mathrm{r}$ from eq. S18 by using eq. S13, eq. S18 is rewritten in the form

$$\Pi b^3 = \phi_\mathrm{r}^2\frac{\partial}{\partial\phi_\mathrm{r}}\left(\frac{f_\mathrm{d}}{\phi_\mathrm{r}}\right). \quad \text{[S19]}$$

The integral constant thus turns out to be the osmotic pressure. By using eqs. S3 - S8, eq. S19 is rewritten in the form

$$\frac{\Pi b^3}{k_\mathrm{B}T} = -(1+\alpha_\mathrm{p})\phi_\mathrm{r} - \log(1-\phi_\mathrm{p}-(1+\alpha_\mathrm{p})\phi_\mathrm{r})$$
$$-\chi(\phi_\mathrm{p}+\alpha_\mathrm{p}\phi_\mathrm{r})^2 - \frac{3(\sigma(r)b^2)^2}{\phi_\mathrm{r}}, \quad \text{[S20]}$$

with

$$\sigma(r) = \frac{\sigma_\mathrm{in}r_\mathrm{in}^2}{r^2}. \quad \text{[S21]}$$

We eliminated $\mu_\mathrm{p}$ in eq. S19 by using eq. S15 to derive the form of eq. S20. So far, we derived the fact that the osmotic pressure is uniform in the DFC layer. The value of the osmotic pressure is determined by taking the limit $r \to r_\mathrm{ex}$ to eq. S18 as

$$\Pi = \Pi_\mathrm{ex} + \frac{2\gamma_\mathrm{ex}}{r_\mathrm{ex}}. \quad \text{[S22]}$$

**C. Relaxation dynamics.** Eq. S13 assumes that the FBRs of nascent pre-rRNAs are relaxed during the transcription of one RNA unit. The time evolution equations of $\alpha_{\rm p}$, $\phi_{\rm p}$, and $\phi_{\rm r}$ are derived by using the Onsager theory. In the steady state, eqs. S11 and S12 still hold because RBPs do not increase or decrease during the transcription. The flux $J$ of nascent pre-rRNA has the form

$$J = -\frac{D_{\rm s}}{k_{\rm B}T}\frac{\partial}{\partial r}(\Pi b^3) \qquad \text{[S23]}$$

where $D_{\rm s}$ is the diffusion constant of a pre-rRNA unit. Because RNA units are added to nascent pre-rRNAs by Pol I at $r = r_{\rm in}$, the condition at the steady state is

$$\frac{4\pi r_{\rm in}\sigma_{\rm in}}{\tau_{\rm s}} = 4\pi r^2 J, \qquad \text{[S24]}$$

where $\tau_{\rm s}$ is the (average) time to transcribe one pre-rRNA unit. The osmotic pressure $\Pi$ satisfies eq. S22 at $r = r_{\rm ex}$ due to the balance of the osmotic pressure, the external pressure, and the capillary force at this position. By solving eq. 29 with this boundary condition, the osmotic pressure is derived as

$$\Pi = \Pi_{\rm ex} + \frac{2\gamma_{\rm ex}}{r_{\rm ex}} + k_{\rm B}T\frac{\sigma_{\rm in}r_{\rm in}^2}{\lambda_{\rm d}^2}\frac{r_{\rm ex}-r}{r_{\rm ex}r}, \qquad \text{[S25]}$$

where $\lambda_{\rm d} = \sqrt{D_{\rm s}\tau_{\rm s}}$ is the diffusion length. The first and second terms of eq. S25 are the same as eq. S22. The RNA units newly added to nascent pre-rRNAs stay at the proximity to the FC-DFC interface until the FBRs of the nascent pre-rRNAs are relaxed. The third term of eq. S25 is the additional pressure generated by the RNA units staying at $r \sim r_{\rm in}$ before the relaxation.

The third term of eq. (S25) scales as

$$\frac{k_{\rm B}T}{b^2}\frac{\sigma_{\rm in}b^2}{\lambda_{\rm d}}\frac{r_{\rm ex}-r_{\rm in}}{\lambda_{\rm d}}\frac{r_{\rm in}}{r_{\rm ex}} \sim 0.3 \text{ Pa}, \qquad \text{[S26]}$$

where the values of parameters used for the estimate are summarized in Table S1. The second term of eq. (S25), $2\gamma_{\rm ex}/r_{\rm ex} \sim$ 2.6 kPa, thus dominates the third term of eq. (S25).

| Parameter | Meaning | Value |
|---|---|---|
| $\sigma_{\rm in}b^2$ | Rescaled surface density of FBRs of nascent pre-rRNAs | 0.1 |
| $r_{\rm ex}$ | External radius of FC-DFC unit | 200 nm |
| $r_{\rm in}$ | Radius of FC | 100 nm |
| $\lambda_{\rm d}$ | Diffusion length | 3 $\mu$m |
| $b$ | Length of RNA unit | 4 nm |
| $T$ | Absolute temperature | 300 K |
| $\gamma_{\rm ex}$ | Surface tension | $2.6 \times 10^{-4}$ N/m |

**Table S1. The values of parameters used in eq. (S26). The diffusion length is estimated in Table 1 in the main article.**

#### S2. Asymptotic solutions for planer geometry

The volume fractions $\phi_{\rm r}$ and $\phi_{\rm p}$ in the layer of nascent pre-rRNAs for the planer geometry are derived by using eqs. S15, S16, and S20 for $\sigma(r) \to \sigma_{\rm in}$. We here derive the asymptotic forms of $\phi_{\rm r}$ and $\phi_{\rm p}$ for each regime.

**A. Good and poor solvent regimes.** For cases in which the interaction parameters $\chi$ is relatively small, the volume fraction $\phi_{\rm r}$ of the FBRs of nascent pre-rRNAs and the volume fraction $\phi_{\rm p}$ of RBPs are relatively small, $\phi_{\rm r} < 1$ and $\phi_{\rm p} < 1$, see Fig. 3 in the main article. With this approximation, eq. S15 has an approximate form

$$\frac{\mu_{\rm p}}{k_{\rm B}T} = \log\phi_{\rm p} \qquad \text{[S27]}$$

Eq. S27 is derived by expanding eq. S15 in a power series of $\phi_{\rm r}$ and $\phi_{\rm p}$ and neglecting higher order terms with respect to these volume fractions. By using eq. S27, the volume fraction of RBPs is derived in the form

$$\phi_{\rm p0} = {\rm e}^{\mu_{\rm p}/(k_{\rm B}T)}. \qquad \text{[S28]}$$

By substituting eq. S28 into eq. S16, the occupancy of the FBRs of nascent pre-rRNAs is derived in the form

$$\alpha_{\rm p0} = \frac{\phi_{\rm p0}{\rm e}^{-(\epsilon-1)/(k_{\rm B}T)}}{1+\phi_{\rm p0}{\rm e}^{-(\epsilon-1)/(k_{\rm B}T)}}. \qquad \text{[S29]}$$

For $\phi_\mathrm{r} < 1$ and $\phi_\mathrm{p} < 1$, eq. S20 has an approximate form

$$\frac{\Pi_\mathrm{ex} b^3}{k_\mathrm{B}T} = -\frac{3(\sigma_\mathrm{in}b^2)^2}{\phi_\mathrm{r}} + (\chi_\mathrm{s} - \chi)\,\alpha_{\mathrm{p0}}^2\phi_\mathrm{r}^2 + \frac{1}{3}(1+\alpha_{\mathrm{p0}})^3\phi_\mathrm{r}^3 \tag{S30}$$

with

$$\chi_\mathrm{s} = \frac{(1+\alpha_{\mathrm{p0}})^2}{2\alpha_{\mathrm{p0}}^2}. \tag{S31}$$

Eq. S30 is derived by expanding eq. S20 in a power series of $\phi_\mathrm{p}$ and $\phi_\mathrm{r}$ and neglecting higher order terms of these volume fractions. We assumed that $\phi_\mathrm{p}$ is very small and neglected even the terms linear to $\phi_\mathrm{p}$. We also used the boundary condition $\Pi = \Pi_\mathrm{ex}$ for the planer geometry, see eq. S22 for $r_\mathrm{ex} \to \infty$. The osmotic pressure has three contributions: the contribution from the entropic elasticity of the RNP complexes (the first term of eq. S30), the contribution from the two-body interaction between the units of RNP complexes (the second term of eq. S30), and the contribution from the three-body interactions between these units (the third term of eq. S30). The two-body interaction between the units of RNA complexes is repulsive for $\chi < \chi_\mathrm{s}$ (the good solvent regime) and is attractive for $\chi > \chi_\mathrm{s}$ (the poor solvent regime). In the good solvent regime, $\chi < \chi_\mathrm{s}$, the first and second terms of eq. S30 dominate the third term of this equation and the volume fraction of the FBRs of nascent pre-rRNA thus has the form

$$\phi_{\mathrm{r0}} = \left(\frac{3(\sigma_\mathrm{in}b^2)^2}{\alpha_{\mathrm{p0}}^2(\chi_\mathrm{s} - \chi)}\right)^{1/3} \tag{S32}$$

if there are not applied pressure to the DFC layer. In the poor solvent regime, $\chi > \chi_\mathrm{s}$, the second and third terms of eq. S30 dominates the third term of this equation and the volume fraction of the FBRs of nascent pre-rRNA has the form

$$\phi_{\mathrm{r0}} = \frac{3\alpha_{\mathrm{p0}}^2(\chi - \chi_\mathrm{s})}{(1+\alpha_{\mathrm{p0}})^3} \tag{S33}$$

Eqs. S32 and S33 correspond to the volume fraction of polymer units for the good and poor solvent conditions, as predicted by the Alexander model, respectively.

**B. Melt regime.** The volume fraction $\phi_\mathrm{r}$ of the FBRs of nascent pre-rRNAs increases with increasing the interaction parameter $\chi$ and, eventually, the RNP complexes occupy most volume of the DFC layer, $\phi_\mathrm{r} \approx 1/(1+\alpha_\mathrm{p})$ (the melt regime), see Fig. 3 in the main article. For simplicity, we here derive the asymptotic forms of the volume fractions $\phi_\mathrm{r}$ and $\phi_\mathrm{p}$ for the case of $\alpha_\mathrm{p} \approx 1$ (which is the case of $\epsilon < -1$). We derive the solution for

$$\phi_\mathrm{r} = \frac{1 - \delta\phi_{\mathrm{rm}}}{2} \tag{S34}$$

$$\alpha_\mathrm{p} = 1 - \delta\alpha_{\mathrm{pm}} \tag{S35}$$

with $\delta\phi_{\mathrm{rm}} \ll 1$, $\delta\alpha_{\mathrm{pm}} \ll 1$, and $\phi_\mathrm{p} \ll 1$. With this approximation, eq. S20 is rewritten in the form

$$\frac{\Pi_\mathrm{ex} b^3}{k_\mathrm{B}T} = -1 - \log(1 - 2\phi_\mathrm{r}) - \frac{1}{4}\chi - 6(\sigma_\mathrm{in}b^2)^2. \tag{S36}$$

Eq. S36 is derived by substituting eqs. S34 and S35, by expanding it in a power series of $\delta\phi_{\mathrm{rm}}$, $\delta\alpha_{\mathrm{pm}}$, and $\phi_\mathrm{p}$. We assumed that $\phi_\mathrm{p}$ is very small and neglected even the terms linear to $\phi_\mathrm{p}$. We also used the boundary condition $\Pi = \Pi_\mathrm{ex}$ for the planer geometry, see eq. S22 for $r_\mathrm{ex} \to \infty$. The volume fraction $\phi_\mathrm{r}$ of the FBRs of nascent pre-rRNAs is derived as

$$\phi_{\mathrm{rm}} = \frac{1}{2}\left(1 - \mathrm{e}^{-1-\frac{1}{4}\chi - \Pi_\mathrm{ex}b^3/(k_\mathrm{B}T)}\right)., \tag{S37}$$

by using eq. S36. To derive eq. S37, we further neglected the fourth term of eq. S36, which is usually smaller than the other terms of eq. S36, see table 1 in the main article.

With the approximation, eqs.S34 and S34, eq. S15 is rewritten in the form

$$\frac{\mu_\mathrm{p}}{k_\mathrm{B}T} = \log\phi_\mathrm{p} - \chi - \log(1 - 2\phi_{\mathrm{rm}}). \tag{S38}$$

The volume fraction of RBPs thus has the form

$$\phi_{\mathrm{pm}} = \mathrm{e}^{\mu_\mathrm{p}/(k_\mathrm{B}T) - 1 + \frac{3}{4}\chi}. \tag{S39}$$

The volume fraction of solvent molecules has the form

$$\phi_\mathrm{s} = 1 - \phi_{\mathrm{pm}} - 2\phi_\mathrm{r} = \mathrm{e}^{-1-\chi/4}\left(1 - \mathrm{e}^{\mu_\mathrm{p}/(k_\mathrm{B}T)+\chi}\right). \tag{S40}$$

The fact that $\phi_\mathrm{s} > 0$ implies that the approximate forms, eqs. S37 and S38, are effective for $\chi + \mu_\mathrm{p}/(k_\mathrm{B}T) < 0$.

**C. DFC regime.** For cases in which the interaction parameter is larger than the negative of the chemical potential of RBPs, $\chi > -\mu_{\rm p}/(k_{\rm B}T)$, most space of the DFC layer is occupied by RBPs (DFC regime), see Fig. 3 in the main article. In this regime, we derive the volume fractions in the form

$$\phi_{\rm p} = 1 - \delta\phi_{\rm pd} \tag{S41}$$

$$\tag{S42}$$

for $\delta\phi_{\rm pd} \ll 1$ and $\phi_{\rm r} \ll 1$. With this approximation, the occupancy of the FBRs of nascent pre-rRNAs has an asymptotic form

$$\alpha_{\rm pd} = \frac{{\rm e}^{-(\epsilon-1)}}{1+{\rm e}^{-(\epsilon-1)}}. \tag{S43}$$

Eq. S43 is derived by substituting eq. S41 into eq. S16, expanding it in a power series of $\delta\phi_{\rm pd}$, and neglecting higher order terms of these parameters.

In the DFC regime, eqs. S15 and S20 has an approximate form

$$\frac{\mu_{\rm p}}{k_{\rm B}T} = -\log(\delta\phi_{\rm p} - (1+\alpha_{\rm pd})\phi_{\rm r}) - 2\chi \tag{S44}$$

$$\frac{\Pi_{\rm ex}b^3}{k_{\rm B}T} = -\log(\delta\phi_{\rm p} - (1+\alpha_{\rm pd})\phi_{\rm r}) - \chi - \frac{3(\sigma_{\rm in}b^2)^2}{\phi_{\rm r}} \tag{S45}$$

where they are derived by substituting eq. S41 into eqs. S15 and S20, expanding them in a power series of $\delta\phi_{\rm pd}$ and $\phi_{\rm r}$, and neglecting the higher order terms of these parameters. We also used the boundary condition $\Pi = \Pi_{\rm ex}$ for the planer geometry, see eq. S22 for $r_{\rm ex} \to \infty$. Eliminating the term $\log(\delta\phi_{\rm p} - (1+\alpha_{\rm pd})\phi_{\rm r})$ in eq. S45 with eq. S44, the relationship between the volume fraction $\phi_{\rm r}$ and the external pressure $\Pi_{\rm ex}$ is derived in the form

$$\frac{\Pi_{\rm ex}b^3}{k_{\rm B}T} = \chi + \frac{\mu_{\rm p}}{k_{\rm B}T} - \frac{3(\sigma_{\rm in}b^2)^2}{\phi_{\rm r}}. \tag{S46}$$

The volume fraction of the FBRs of nascent pre-rRNAs thus has the form

$$\phi_{\rm rd} = \frac{3(\sigma_{\rm in}b^2)^2}{\chi + (\mu_{\rm p} - \Pi_{\rm ex}b^3)/(k_{\rm B}T)}. \tag{S47}$$

#### S3. Asymptotic form of spherical geometry

The local volume fractions of the FBRs of nascent pre-rRNA $\phi_{\rm r}(r)$ and RBPs $\phi_{\rm p}(r)$ are derived by using eqs. S15, S16, S17, and S22 for the spherical geometry. We derive the volume fractions, the free energy $F$ of the system, and the radius $r_{\rm in}$ of FCs for $\alpha_{\rm p} \approx 1$ and $\chi > -\mu_{\rm p}/(k_{\rm B}T)$. The approximation, $\alpha_{\rm p} \approx 1$, is effective for $\epsilon - 1 - \mu_{\rm p}/(k_{\rm B}T) \ll -1$, where RBPs are bound to the FBRs of pre-rRNA even in a dilute solution. In this case, the layer composed of the FBRs of nascent pre-rRNAs can be a uniform DFC layer, a uniform melt layer, and a double layer of DFC and melt regions.

**A. Uniform DFC layer.** In the DFC regime, most volume of the layer composed of the FBRs of nascent pre-rRNAs is occupied by RBPs. We thus derive the solution of eqs. S13, S15, S16, and S22 in the form of

$$\phi_{\rm p}(r) = 1 - \delta\phi_{\rm p}(r), \tag{S48}$$

where $\delta\phi_{\rm p} \ll 1$ and $\phi_{\rm r} \ll 1$. In this case, eq. S15 has an approximate form

$$\frac{\mu_{\rm p}}{k_{\rm B}T} = -\log(\delta\phi_{\rm p}(r) - 2\phi_{\rm r}(r)) - 2\chi. \tag{S49}$$

Eq. S49 is derived by substituting eq. S48, expanding it in a power series of $\delta\phi_{\rm p}(r)$ and $\phi_{\rm r}$, and neglecting the higher order terms with respect to $\delta\phi_{\rm p}(r)$ and $\phi_{\rm r}$. By using the same expansion and approximation, eq. S17 is rewritten in the form

$$\begin{aligned}\frac{\mu_{\rm r}}{k_{\rm B}T} &= -\frac{3}{2}\frac{(\sigma_{\rm in}b^2)^2}{\phi_{\rm r}^2(r)}\frac{r_{\rm in}^4}{r^4} - 2\log(\delta\phi_{\rm p} - 2\phi_{\rm r}) - 2 - 2\chi + \epsilon - \frac{\mu_{\rm p}}{k_{\rm B}T} \\ &= -\frac{3}{2}\frac{(\sigma_{\rm in}b^2)^2}{\phi_{\rm r}^2(r)}\frac{r_{\rm in}^4}{r^4} + \frac{\mu_{\rm p}}{k_{\rm B}T} + 2\chi + \epsilon - 2.\end{aligned} \tag{S50}$$

The last form of eq. S50 is derived by eliminating $\log(\delta\phi_{\rm p} - 2\phi_{\rm r})$ by using eq. S49. By solving eq. S50 with respect to $\phi_{\rm r}(r)$, the volume fraction $\phi_{\rm r}(r)$ of the FBRs of nascent pre-rRNAs is derived as

$$\phi_{\rm r}(r) = \sqrt{\frac{3}{2}}\frac{\sigma_{\rm in}b^2}{\sqrt{\Delta\mu_{\rm r}/(k_{\rm B}T)}}\frac{r_{\rm in}^2}{r^2}, \tag{S51}$$

where we introduced a parameter

$$\frac{\Delta\mu_{\rm r}}{k_{\rm B}T} = \frac{\mu_{\rm p}}{k_{\rm B}T} + 2\chi + \epsilon - 2 - \frac{\mu_{\rm r}}{k_{\rm B}T} \quad \text{[S52]}$$

to simplify the expression of eq. S51.

The osmotic pressure has an approximate form

$$\begin{aligned} \frac{\Pi b^3}{k_{\rm B}T} &= -\log(\delta\phi_{\rm p}(r) - 2\phi_{\rm r}(r)) - \chi - \frac{3(\sigma_{\rm in}b^2)^2}{\phi_{\rm r}(r_{\rm ex})}\frac{r_{\rm in}^4}{r_{\rm ex}^4} \\ &= \chi + \frac{\mu_{\rm p}}{k_{\rm B}T} - \frac{3(\sigma_{\rm in}b^2)^2}{\phi_{\rm r}(r)}\frac{r_{\rm in}^4}{r^4}, \end{aligned} \quad \text{[S53]}$$

where it is derived by substituting eq. S48 into eq. S20, expanding it in a power series of $\delta\phi_{\rm p}(r)$ and $\phi_{\rm r}(r)$, and neglecting the higher order terms with respect to $\delta\phi_{\rm p}(r)$ and $\phi_{\rm r}(r)$. The last form of eq. S53 is derived by eliminating $\log(\delta\phi_{\rm p}(r) - 2\phi_{\rm r}(r))$ by using eq. S50. By using eq. S22, the volume fraction $\phi_{\rm rex}$ $(= \phi_{\rm r}(r_{\rm ex}))$ of the FBRs of nascent pre-rRNAs at $r = r_{\rm ex}$ is derived as

$$\phi_{\rm rex} = \frac{3(\sigma_{\rm in}b^2)^2}{\chi + (\mu_{\rm p} - \Pi_{\rm ex})/(k_{\rm B}T) - 2\gamma_{\rm p}b^3/r_{\rm ex}}\frac{r_{\rm in}^4}{r_{\rm ex}^4}. \quad \text{[S54]}$$

By using the boundary condition, eq. S54, the local volume fraction $\phi_{\rm r}(r)$ of the FBRs of nascent pre-rRNA has the form

$$\phi_{\rm r}(r) = \phi_{\rm rex}\frac{r_{\rm ex}^2}{r^2}. \quad \text{[S55]}$$

The chemical potential $\mu_{\rm r}$ has the form

$$\frac{\mu_{\rm r}}{k_{\rm B}T} = \frac{\mu_{\rm p}}{k_{\rm B}T} + 2\chi + \epsilon - 2 - \frac{1}{6}\frac{1}{(\sigma_{\rm in}b^2)^2}\left(\chi + \frac{\mu_{\rm p} - \Pi_{\rm ex}}{k_{\rm B}T} - \frac{2\gamma_{\rm p}b^3}{r_{\rm ex}}\right)\frac{r_{\rm ex}^4}{r_{\rm in}^4}. \quad \text{[S56]}$$

Eqs. S54, S55, and S56 are effective for cases in which the layer of the FBRs of nascent pre-rRNAs is a double layer of the DFC and melt phases if the DFC layer interfaces with GC at $r = r_{\rm ex}$.

The external radius $r_{\rm ex}$ is derived by using the relationship

$$\frac{r_{\rm in}^2}{r_{\rm ex}^2}\left(\frac{r_{\rm ex}}{r_{\rm in}} - 1\right) = \frac{N_{\rm r}b}{3(\sigma_{\rm in}b^2)r_{\rm in}}\left(\chi + \frac{\mu_{\rm p} - \Pi_{\rm ex}}{k_{\rm B}T} - \frac{2\gamma_{\rm p}b^3}{r_{\rm ex}}\right), \quad \text{[S57]}$$

where it is derived by substituting eq. S55 (and eq. S54) into eq. S4. Indeed, eq. S57 is a quadratic equation and the ratio $r_{\rm ex}/r_{\rm in}$ can be derived analytically.

The free energy density of the layer of the FBRs of nascent pre-rRNAs has an approximate form

$$\begin{aligned} \frac{f_{\rm d}}{k_{\rm B}t} &= \frac{3}{2}\frac{(\sigma_{\rm in}b^2)^2}{\phi_{\rm r}(r)}\frac{r_{\rm in}^4}{r^4} - \left(\chi + \frac{\mu_{\rm p} - \Pi_{\rm ex}}{k_{\rm B}T}\right) \\ &= \frac{1}{2}\left(\chi + \frac{\mu_{\rm p} - \Pi_{\rm ex}}{k_{\rm B}T} - 2\gamma_{\rm p}b^3/r_{\rm ex}\right)\frac{r_{\rm ex}^2}{r^2} - \left(\chi + \frac{\mu_{\rm p} - \Pi_{\rm ex}}{k_{\rm B}T}\right). \end{aligned} \quad \text{[S58]}$$

Eq. S58 is derived by substituting eqs. S48, expanding it in a a power series of $\delta\phi_{\rm p}$ and $\phi_{\rm r}$, and neglecting the higher order terms of $\delta\phi_{\rm p}$ and $\phi_{\rm r}$. We also used eq. S55 to derive the last form of eq. S58. By using eq. S58, the free energy of the system is derived as

$$\begin{aligned} \frac{Fb^3}{3V_{\rm m}k_{\rm B}T} &= \frac{1}{6}\left(\chi + \frac{\mu_{\rm p} - \Pi_{\rm ex}}{k_{\rm B}T}\right)\left[\left(\frac{r_{\rm ex}}{r_{\rm in}}\right)^3 - 3\left(\frac{r_{\rm ex}}{r_{\rm in}}\right)^2 + 2\right] + \frac{\gamma_{\rm p}b^3}{k_{\rm B}Tr_{\rm in}}\left(1 + \frac{r_{\rm ex}}{r_{\rm in}}\right) \\ &\quad + \zeta\left(\epsilon + 2\chi + \frac{\mu_{\rm p}}{k_{\rm B}T} - 2\right), \end{aligned} \quad \text{[S59]}$$

where we used

$$\zeta = N_{\rm r}\frac{\sigma_{\rm in}b^3}{r_{\rm in}}. \quad \text{[S60]}$$

$\zeta$ is proportional to the transcription rate. The free energy, eq. S59, depends on the size of FCs via the ratio $r_{\rm ex}/r_{\rm in}$ and $\sigma_{\rm in}(\propto r_{\rm in})$, see also eq. S57.

For $\gamma_{\rm p}b^3/r_{\rm in} \ll \chi + (\mu_{\rm p} - \Pi_{\rm ex})/(k_{\rm B}T)$, the first term of eq. S59 dominates the second term of this equation. In this case, the first derivative of the free energy with respect to the raio $r_{\rm ex}$/in is zero at the minimum of the free energy,

$$\frac{\partial}{\partial(r_{\rm ex}/r_{\rm in})}\left(\frac{Fb^3}{3V_{\rm m}k_{\rm B}T}\right) \simeq \frac{1}{2}\left(\chi + \frac{\mu_{\rm p} - \Pi_{\rm ex}}{k_{\rm B}T}\right)\frac{r_{\rm ex}}{r_{\rm in}}\left(\frac{r_{\rm ex}}{r_{\rm in}} - 2\right) = 0, \quad \text{[S61]}$$

This leads to

$$\frac{r_{\rm ex}}{r_{\rm in}} = 2. \qquad \text{[S62]}$$

By substituting eq. S62 into eq. S57, the radius of FCs is derived as

$$\frac{r_{\rm in}}{N_{\rm r} b} = \frac{\sigma_{\rm in} b^2}{\zeta} = \frac{2}{\sqrt{3}}\left(\chi + \frac{\mu_{\rm p} - \Pi_{\rm ex}}{k_{\rm B}T}\right)^{1/2}\zeta^{-1/2}, \qquad \text{[S63]}$$

implying that the radius of FCs is proportional to the inverse of the square root of the transcription rate.

**B. Uniform melt layer.** For cases in which the FBRs of pre-rRNAs form a uniform melt layer, the volume fraction $\phi_{\rm r}$ of the FBRs of pre-rRNAs is $\approx 1/2$. We therefore derive the volume fractions in the forms

$$\phi_{\rm r} = \frac{1}{2}(1 - \delta\phi_{\rm r}), \qquad \text{[S64]}$$

where $\phi_{\rm p} \ll 1$ and $\delta\phi_{\rm r} \ll 1$. We substitute eq. S64 into eqs. S15, S20, S17, expanding these equations in a power series of $\phi_{\rm p}$ and $\delta\phi_{\rm r}$, and neglecting the higher order terms with respect to $\phi_{\rm p}$ and $\delta\phi_{\rm r}$,

$$\frac{\mu_{\rm p}}{k_{\rm B}T} = \log\phi_{\rm p} - \log(\delta\phi_{\rm r} - \delta\phi_{\rm p}) - \chi \qquad \text{[S65]}$$

$$\frac{\mu_{\rm r}}{k_{\rm B}T} = -6(\sigma(r)b^2)^2 - 2\log(\delta\phi_{\rm r} - \delta\phi_{\rm p}) - 2 - \chi + \epsilon - \frac{\mu_{\rm p}}{k_{\rm B}T} \qquad \text{[S66]}$$

$$\frac{\Pi b^3}{k_{\rm B}T} = -\log(\delta\phi_{\rm r} - \phi_{\rm p}) - 1 - \frac{1}{4}\chi - 6(\sigma(r)b^2)^2. \qquad \text{[S67]}$$

The osmotic pressure $\Pi$ is derived by eliminating $\log(\delta\phi_{\rm r} - \phi_{\rm p})$ in eq. S67 by using eq. S66. This leads to the form

$$\frac{\Pi b^3}{k_{\rm B}T} = \frac{1}{2}\left(\frac{\mu_{\rm r} + \mu_{\rm p}}{k_{\rm B}T} - \epsilon\right) + \frac{1}{4}\chi - 3(\sigma(r)b^2)^2. \qquad \text{[S68]}$$

By substituting eq. S66 into eq. S4, the ratio $r_{\rm ex}/r_{\rm in}$ is derived as

$$\frac{r_{\rm ex}}{r_{\rm in}} = (1 + 6\zeta)^{1/3}, \qquad \text{[S69]}$$

where $\zeta$ is given by eq. S60. The ratio $r_{\rm ex}/r_{\rm in}$ thus does not depend on the radius of FCs. The free energy has an approximate form

$$\frac{Fb^3}{3V_{\rm m}k_{\rm B}T} = -\frac{1}{6}\left(\frac{1}{2}\chi + \frac{\mu_{\rm p}}{k_{\rm B}T}\right)\left(\frac{r_{\rm ex}^3}{r_{\rm in}^3} - 1\right) + 3(\sigma_{\rm in}b^2)^2\left(1 - \frac{r_{\rm in}}{r_{\rm ex}}\right) + \frac{1}{2}\frac{\gamma_{\rm p}b^3}{k_{\rm B}Tr_{\rm in}}\left(1 + \frac{r_{\rm ex}^2}{r_{\rm in}^2}\right), \qquad \text{[S70]}$$

where it is derived by using $\phi_{\rm r} \approx 1/2$ and $\phi_{\rm p} \simeq 1 - 2\phi_{\rm r} \ll 1$ to eq. S1. The derivative of the free energy with respect to $\sigma_{\rm in}b^2$ is zero at the minimum of the free energy,

$$\frac{\partial}{\partial(\sigma_{\rm in}b^2)}\left(\frac{Fb^3}{3V_{\rm m}k_{\rm B}T}\right) = 6(\sigma_{\rm in}b^2)\left(1 - \frac{r_{\rm in}}{r_{\rm ex}}\right) - \frac{1}{2}\frac{\gamma_{\rm p}b^2\zeta}{N_{\rm r}k_{\rm B}T}\frac{1}{(\sigma_{\rm in}b^2)^2}\left(1 + \frac{r_{\rm ex}^2}{r_{\rm in}^2}\right) = 0. \qquad \text{[S71]}$$

The radius of FCs thus has the form

$$\frac{r_{\rm in}}{N_{\rm r}b} = \left(\frac{\gamma_{\rm p}b^2}{12N_{\rm r}k_{\rm B}T}\right)^{1/3}\zeta^{-2/3}\left(\frac{1 + r_{\rm ex}^2/r_{\rm in}^2}{1 - r_{\rm in}/r_{\rm ex}}\right)^{1/3}, \qquad \text{[S72]}$$

see eq. S69 for the dependence of $r_{\rm ex}/r_{\rm in}$ on $\zeta$. For $\zeta \ll 1$, eq. S72 has an approximate form

$$\frac{r_{\rm in}}{N_{\rm r}b} \simeq \left(\frac{\gamma_{\rm p}b^2}{12N_{\rm r}k_{\rm B}T}\right)^{1/3}\zeta^{-1}. \qquad \text{[S73]}$$

**C. Double layer of DFC and melt.** In some cases, the layer occupied by the FBRs of nascent pre-rRNAs is composed of a double layer of a melt phase and a DFC phase. The chemical potentials, $\mu_{\rm p}$ and $\mu_{\rm r}$, and the osmotic pressure $\Pi$ is continuous at the interface between the two layers. The osmotic pressure increases with increasing the radial coordinate $r$ in the DFC layer, see eq. S53, while the osmotic pressure is almost constant in the melt layer, see eq. S68 (where the third term of eq. S68 is smaller than the other terms in this equation and is neglected). This implies that the DFC layer is at the interface with the GC and the melt layer at the interface with the FC. The osmotic pressure in the DFC layer has the form

$$\frac{\Pi b^3}{k_{\rm B}T} = \chi + \frac{\mu_{\rm p}}{k_{\rm B}T} - \left(\chi + \frac{\mu_{\rm p} - \Pi_{\rm ex} b^3}{k_{\rm B}T} - \frac{2\gamma_{\rm ex} b^3}{k_{\rm B}T r_{\rm ex}}\right)\frac{r_{\rm ex}^2}{r^2}, \qquad [S74]$$

where it is derived by substituting eq. S55 into eq. S53. The osmotic pressure in the melt layer has the form

$$\frac{\Pi b^3}{k_{\rm B}T} = -\frac{1}{12(\sigma_{\rm in} b^2)^2}\left(\chi + \frac{\mu_{\rm p} - \Pi_{\rm ex} b^3}{k_{\rm B}T} - \frac{2\gamma_{\rm ex} b^3}{k_{\rm B}T r_{\rm ex}}\right)^2 \frac{r_{\rm ex}^4}{r_{\rm in}^4} + \frac{\mu_{\rm p}}{k_{\rm B}T} + \frac{5}{4}\chi - 1, \qquad [S75]$$

where it is derived by substituting the chemical potential

$$\frac{\mu_{\rm r}}{k_{\rm B}T} = -\frac{3}{2}\frac{(\sigma_{\rm in} b^2)^2}{\phi_{\rm rex}^2}\frac{r_{\rm in}^4}{r_{\rm ex}^4} + \frac{\mu_{\rm p}}{k_{\rm B}T} + 2\chi + \epsilon - 2. \qquad [S76]$$

in eq. S68 and neglected the third term of eq. S68. Eq. S76 is derived by substituting eq. S55 into eq. S50 and is effective for cases in which the DFC layer interfaces with the GC (because eq. S68 is imposed at $r = r_{\rm ex}$). The radical coordinate $r_{\rm i}$ at the interface between the DFC and melt layer is derived by equating eqs. S74 and S75,

$$\frac{r_{\rm i}^2}{r_{\rm in}^2} = \frac{\left(\chi + \frac{\mu_{\rm p} - \Pi_{\rm ex} b^3}{k_{\rm B}T} - \frac{2\gamma_{\rm ex} b^3}{k_{\rm B}T r_{\rm ex}}\right)\frac{r_{\rm ex}^2}{r_{\rm in}^2}}{1 - \frac{1}{4}\chi + \frac{1}{12(\sigma_{\rm in} b^2)^2}\left(\chi + \frac{\mu_{\rm p} - \Pi_{\rm ex} b^3}{k_{\rm B}T} - \frac{2\gamma_{\rm ex} b^3}{k_{\rm B}T r_{\rm ex}}\right)^2 \frac{r_{\rm ex}^4}{r_{\rm in}^4}}. \qquad [S77]$$

The volume fraction $\phi_{\rm r}$ of the units of nascent pre-rRNAs is given by eq. S55 in the DFC layer ($r_{\rm i} < r < r_{\rm ex}$) and is $\approx 1/2$ in the melt layer ($r_{\rm in} < r < r_{\rm i}$),

$$\phi_{\rm r}(r) = \begin{cases} \frac{1}{2} & (r_{\rm in} < r < r_{\rm i}) \\ \phi_{\rm rex}\frac{r_{\rm ex}^2}{r^2} & (r_{\rm i} < r < r_{\rm ex}) \end{cases} \qquad [S78]$$

A relationship

$$\frac{3(\sigma_{\rm in} b^2)^2}{\chi + \frac{\mu_{\rm p} - \Pi_{\rm ex} b^3}{k_{\rm B}T} - \frac{2\gamma_{\rm ex} b^3}{k_{\rm B}T r_{\rm ex}}}\frac{r_{\rm in}^2}{r_{\rm ex}^2}\left(\frac{r_{\rm ex}}{r_{\rm in}} - \frac{r_{\rm i}}{r_{\rm in}}\right) + \frac{1}{6}\left(\frac{r_{\rm i}^3}{r_{\rm in}^3} - 1\right) = \zeta. \qquad [S79]$$

is derived by substituting eq. S78 into eq. S4. By using eqs. S77 and S79, one can derive the ratio $r_{\rm i}/r_{\rm in}$ and $r_{\rm ex}/r_{\rm in}$ as a function of $\sigma_{\rm in} b^2$ ($\propto r_{\rm in}$). The free energy of the system is derived as

$$\begin{aligned} \frac{F b^3}{3 k_{\rm B}T V_{\rm m}} = {} & \frac{1}{6}\left(\chi + \frac{\mu_{\rm p} - \Pi_{\rm ex} b^3}{k_{\rm B}T}\right)\left(\frac{r_{\rm ex}^3}{r_{\rm in}^3} - 3\frac{r_{\rm ex}^2}{r_{\rm in}^2}\frac{r_{\rm i}}{r_{\rm in}} + 2\frac{r_{\rm i}^3}{r_{\rm in}^3}\right) + \frac{\gamma_{\rm p} b^3}{k_{\rm B}T r_{\rm in}}\left(\frac{1}{2} + \frac{r_{\rm ex} r_{\rm i}}{r_{\rm in}^2}\right) + \frac{\gamma_{\rm p} b^3}{k_{\rm B}T r_{\rm in}}\frac{1}{2}\frac{r_{\rm i}^2}{r_{\rm in}^2} \\ & -\frac{1}{6}\left(\frac{1}{2}\chi + \frac{\mu_{\rm p}}{k_{\rm B}T} - \frac{2\Pi_{\rm ex} b^3}{k_{\rm B}T} - \epsilon\right)\left(\frac{r_{\rm i}^3}{r_{\rm in}^3} - 1\right) \\ & + \frac{3(\sigma_{\rm in} b^2)^2}{\chi + \frac{\mu_{\rm p} - \Pi_{\rm ex} b^3}{k_{\rm B}T} - \frac{2\gamma_{\rm ex} b^3}{k_{\rm B}T r_{\rm ex}}}\left(\epsilon + 2\chi + \frac{\mu_{\rm p}}{k_{\rm B}T} - 2\right)\frac{r_{\rm in}^2}{r_{\rm ex}^2}\left(\frac{r_{\rm ex}}{r_{\rm in}} - \frac{r_{\rm i}}{r_{\rm in}}\right). \end{aligned} \qquad [S80]$$

#### S4. Contribution of DNA to FC size

Not only the terminal regions of nascent pre-rRNAs, but also rDNAs are localized at the surfaces of FCs. One may think that rDNAs also contribute to the suppression of the growth of FCs. The rDNA is bound to the surface of FCs and thus bent to the curvature of the surface. With these contributions, the free energy has the form

$$F = \frac{3\gamma_{\rm in} V_{\rm m}}{r_{\rm in}} + k_{\rm B}T\frac{l_{\rm DNA} L}{2 r_{\rm in}^2} - k_{\rm B}T\epsilon_{\rm DNA} \qquad [S81]$$

This free energy is composed of the surface free energy due to the surface tension $\gamma_{\rm in}$ between FC and DFC (the first term), the elastic free energy due to the bending of rDNA (the second term), and is the free energy due to the binding of rDNA to the surface of FCs (the third term). $k_{\rm B}T\epsilon_{\rm DNA}$ is the free energy due to the binding of rDNA, $l_{\rm DNA}$ ($\approx 50$ nm) is the persistence length of DNA, and $L_{\rm DNA}$ is the total length of rDNA bound at the surface of FCs. Eq. S81 is derived by using the fact that the sum of the volume $V_{\rm m}$ of FCs is constant. Both the surface free energy and the bending free energy of rDNA decreases with increasing the radius $r_{\rm in}$ of FCs (see the first and second terms in eq. S81), while the binding free energy is constant (see the third term in eq. S81), implying that rDNA itself does not have a function to suppress the fusion of FCs. If we assume that the radius of FCs is $r_{\rm in} \sim 100$ nm, the first term is estimated as $9 \times 10^{-19}$ J (by using $\gamma \sim 1 \times 10^{-6}$ N/m), while the second term is estimated as $6 \times 10^{-20}$J (by using the fact that there are typically 10 FCs per nucleolus (1) and the persistence length of DNA is $l_{\rm DNA} \approx 50$ nm in the physiological salt concentration (2)), see also Table 2. This estimate implies that the first term of eq. S81 dominates the second term of eq. S81 for the FC radius $r_{\rm in}$ treated in our theory, $r_in \leq 100$ nm and experiments. The above estimate does not take into account the fact that a stretch of DNA can be associated with multiple FCs. It is well-known in the colloid science that the polymers bridging between liquid droplets rather drive the coalescence of these droplets due to their entropic elasticity (3).

#### S5. Elastic free energy of nascent RNA transcripts

In the main article, we use an extension of the theory of polymer brush to treat nascent pre-rRNA transcripts. The Alexander approximation assumes that the volume fraction of nascent pre-rRNA units is uniform for nascent pre-rRNA transcripts on a planer surface (4, 5). With this approximation, the free energy per unit area of nascent pre-rRNA transcripts on a planer surface has the form

$$\frac{F_{\rm bru}}{k_{\rm B}T} = \frac{3}{2}\frac{\sigma h^2}{N_{\rm r}b^2} + v\sigma N_{\rm r}\frac{\sigma N_{\rm r}}{h}. \quad [S82]$$

This free energy is a function of the height $h$ of the nascent pre-rRNA transcripts. The first term of eq. S82 is the elastic free energy of nascent pre-rRNA transcripts and the second term of eq. S82 is the free energy due to the interactions between nascent pre-rRNA units. $k_{\rm B}$ is the Boltzmann constant and $T$ is the absolute temperature. $N_{\rm r}$ is the number of units in a nascent pre-rRNA transcript and $b$ is the length of each unit. $\sigma$ is the surface density of nascent pre-rRNA transcripts. $v$ is the excluded volume that represents the magnitude of the interactions between nascent pre-rRNA transcripts and has a relationship $v = b^3(2-\chi)$ with the interaction parameter $\chi$ introduced in eq. 12 in the main article (the difference from the usual relationship $b^3(1-2\chi)/2$ is due to the volume of RBPs bound to nascent pre-rRNA transcripts. Eq. S82 is rewritten in the form

$$\frac{F_{\rm bru}}{k_{\rm B}T} = \frac{3}{2}\frac{b^4\sigma^3 N_{\rm r}}{\phi_{\rm r}^2} + \frac{v}{b^3}\sigma N_{\rm r}\phi_{\rm r} \quad [S83]$$

by using the form of the volume fraction of nascent pre-rRNA units

$$\phi_{\rm r} = \frac{b^3\sigma N_{\rm r}}{h}. \quad [S84]$$

Minimizing eq. S82 with respect to $h$ leads to

$$h = N_{\rm r}b\left(\frac{\sigma v}{3b}\right)^{1/3}. \quad [S85]$$

The volume fraction of nascent pre-rRNA units is derived as

$$\phi_{\rm r} = \left(\frac{3\sigma^2 b^7}{v}\right)^{1/3} \quad [S86]$$

by substituting eq. S85 into eq. S84 or by minimizing eq. S83 with respect to $\phi_{\rm r}$.

Eq. S82 is rewritten in the following form

$$\frac{F_{\rm bru}}{k_{\rm B}T} = \frac{\sigma N_{\rm r}}{g}\left[\frac{3}{2}\frac{\xi^2}{gb^2} + vg\frac{g}{\xi^3}\right]. \quad [S87]$$

With this expression, we divide nascent pre-rRNA transcripts into blobs of size $\xi$ ($\equiv \sigma^{-1/2}$), which the average distance between grafting points (to which nascent pre-rRNA transcripts are end-grafted) at the surface. $g$ is the number of nascent pre-rRNA units in each blob. The first term in the square bracket in eq. S87 is the elastic free energy of the subchain in a blob and the second term in the square bracket is the free energy due to the interactions between nascent pre-rRNA units in the blob. The prefactor $N_{\rm r}/g$ is the number of blobs in each nascent pre-rRNA transcript. Eq. S87 returns to eq. S83 by using the form of the local volume fraction of nascent pre-rRNA units

$$\phi_{\rm r} = \frac{b^3 g}{R^3}. \quad [S88]$$

Now we extend the above formalism to nascent pre-rRNA transcripts on a spherical surface, see Fig. 1**b** in the main article. The size of each blob is

$$\xi = \frac{r}{\sigma^{1/2}r_{\rm in}} \quad [S89]$$

due to the fact that the spherical surface at the distance $r$ from the center of this surface is occupied by $4\pi r_{\rm in}^2\sigma$ transcripts,

$$4\pi r^2 = (4\pi r_{\rm in}^2\sigma)\xi^2. \qquad [S90]$$

The number of blobs in the spherical shell of thickness $dr$ at the distance $r$ from the center is $4\pi r^2 dr/\xi^3$. Eq. S87 is extended to

$$\frac{F_{\rm bru}}{k_{\rm B}T} = \int_{r_{\rm in}}^{r_{\rm ex}} \frac{4\pi r^2 dr}{\xi^3}\left[\frac{3}{2}\frac{\xi^2}{gb^2} + vg\frac{g}{\xi^3}\right], \qquad [S91]$$

where it is along with the spirit of the Daoud-Cotton theory (6). Eq. S91 is rewritten in the form

$$\frac{F_{\rm bru}}{k_{\rm B}T} = \int_{r_{\rm in}}^{r_{\rm ex}} \frac{4\pi r^2 dr}{b^3}\left[\frac{3}{2}\frac{b^4\sigma^2 r_{\rm in}^4}{r^4}\frac{1}{\phi_{\rm r}} + \frac{v}{b^3}\phi_{\rm r}^2\right] \qquad [S92]$$

by using eq. S88 and $R = \sigma^{-1/2}$. The first term in the square bracket of eq. S92 is the elastic free energy of the chain subsection in a blob of nascent pre-rRNA transcripts and the second term in the square bracket of eq. S92 is the free energy due to the interactions between nascent pre-rRNA units in the blob. We thus used the first term of eq. S92 for the elastic free energy, see eq. 10 in the main article.

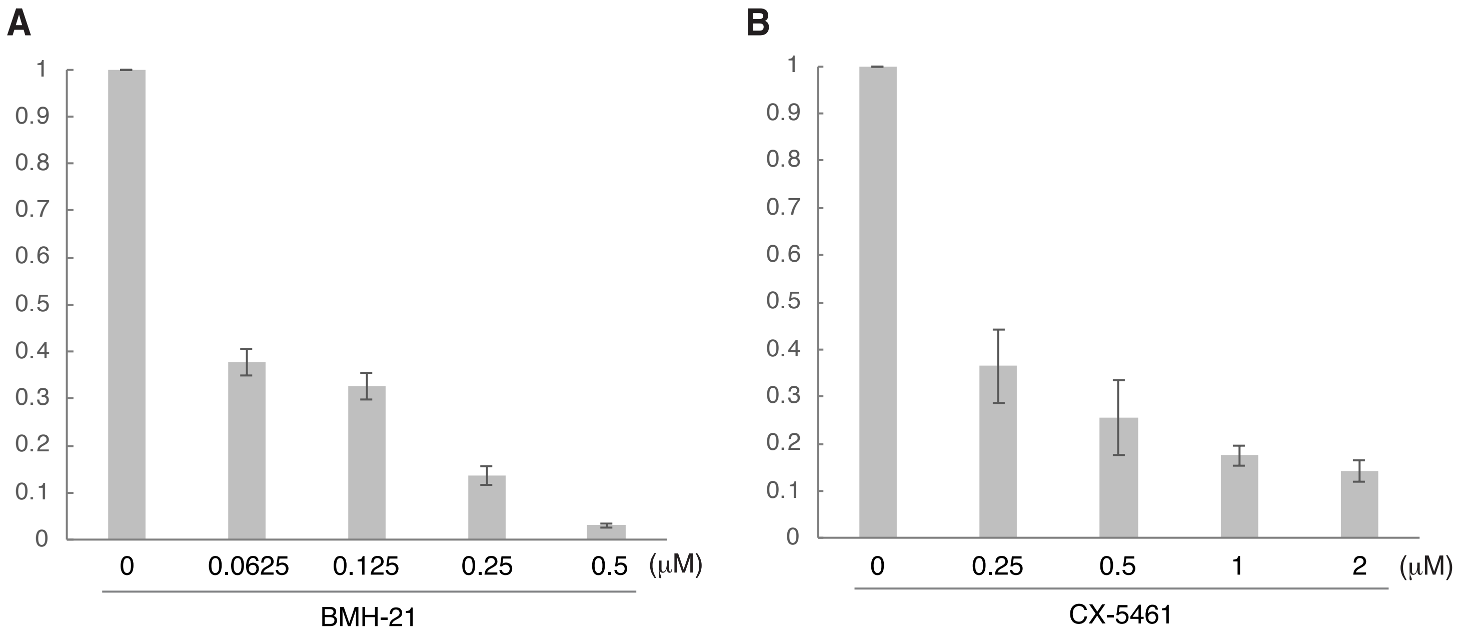


**Figure S1 BMH-21 and CX-5461 treatment reduce pre-rRNA expression levels in a dose-dependent manner.**

(*A*) Quantification of pre-rRNA expression levels by RT-qPCR in BMH-21-untreated and -treated conditions. Data are represented as mean ± SD (*n* =3). (*B*) Quantification of pre-rRNA expression levels by RT-qPCR in CX-5461-untreated and -treated conditions. Data are represented as mean ± SD (*n* =3).

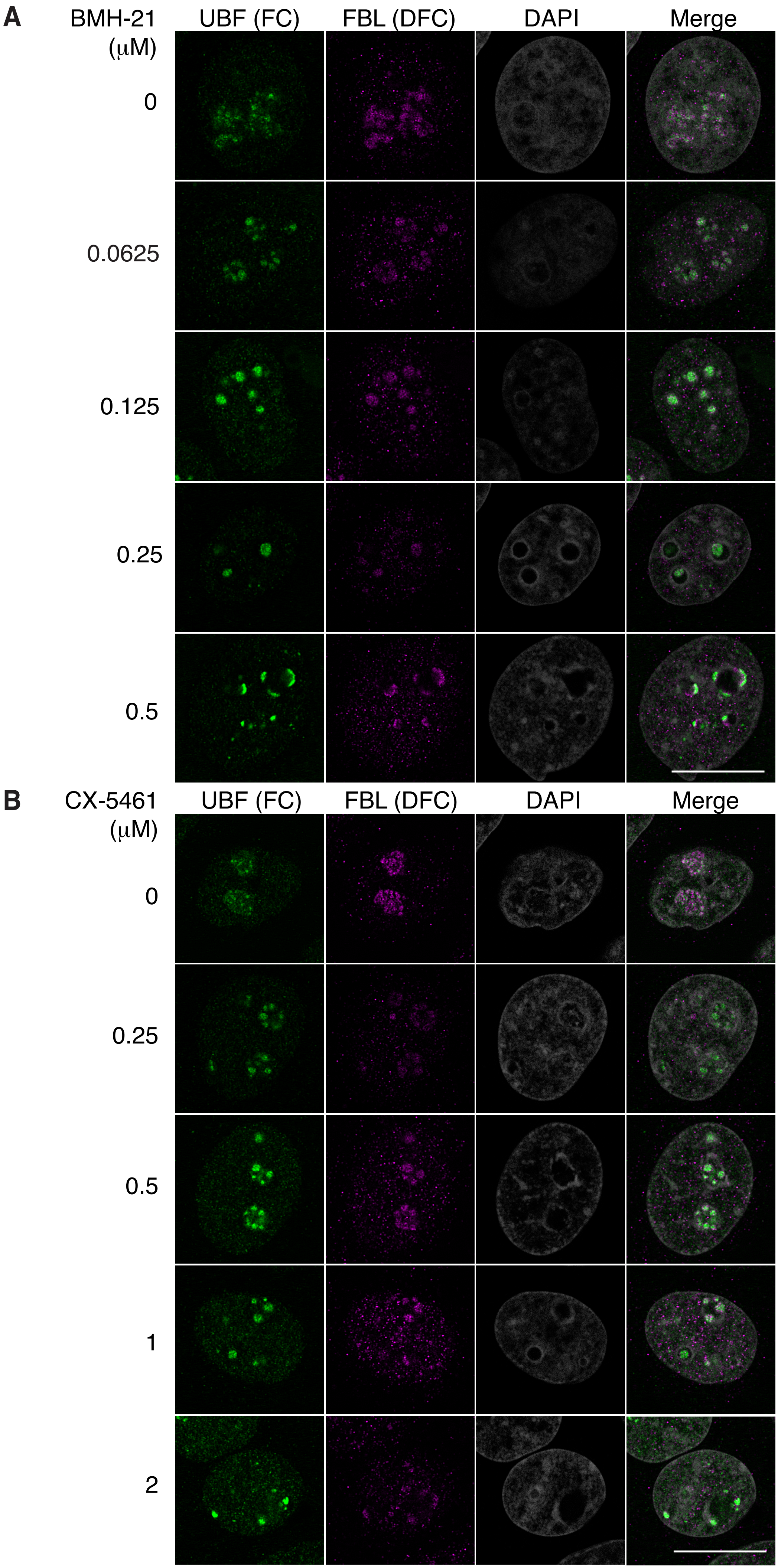
A
BMH-21 (μM)
UBF (FC)
FBL (DFC)
DAPI
Merge
0
0.0625
0.125
0.25
0.5
B
CX-5461 (μM)
UBF (FC)
FBL (DFC)
DAPI
Merge
0
0.25
0.5
1
2

**Figure S2 FCs and DFCs in the BMH-21 and CX-5461-untreated and -treated cells.** (*A* and *B*) Immunofluorescence of UBF (FC) and FBL (DFC) in HeLa cells with or without BMH-21 (*A*) or CX-5461 (*B*) treatments. Scale bar, 10 µm.

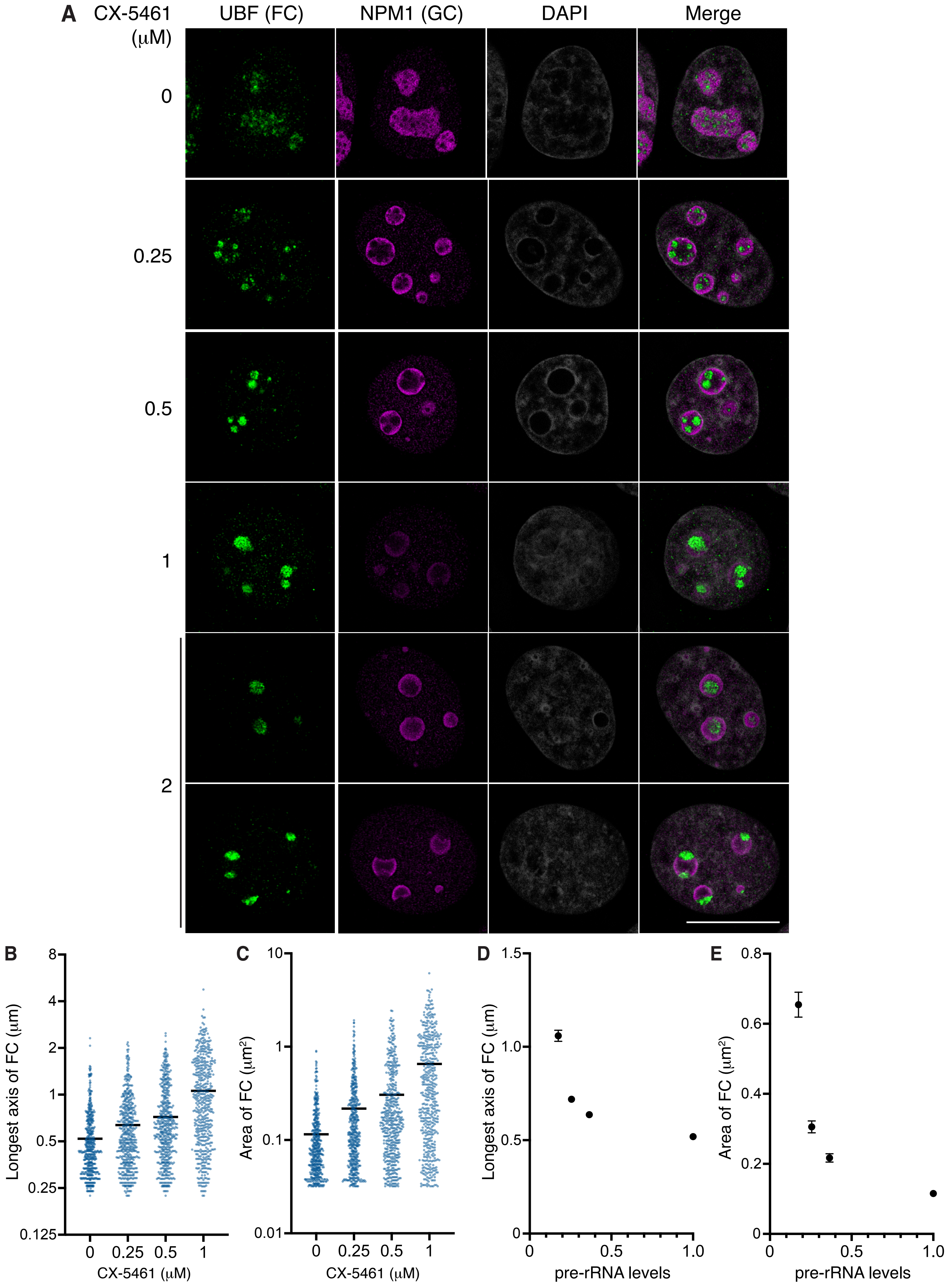

A
CX-5461 (μM)
UBF (FC)
NPM1 (GC)
DAPI
Merge
0
0.25
0.5
1
2
B
Longest axis of FC (μm)
8
4
2
1
0.5
0.25
0.125
0
0.25
0.5
1
CX-5461 (μM)
C
Area of FC (μm²)
10
1
0.1
0.01
0
0.25
0.5
1
CX-5461 (μM)
D
Longest axis of FC (μm)
1.5
1.0
0.5
0
0
0.5
1.0
pre-rRNA levels
E
Area of FC (μm²)
0.8
0.6
0.4
0.2
0
0
0.5
1.0
pre-rRNA levels

**Figure S3  Mild Pol I inhibition by CX-5461 increases the size of FCs.**

(*A*) Immunofluorescence of UBF (FC) and NPM1 (GC) in HeLa cells with or without CX-5461 treatments. In the cells treated with 2 μM CX -5461, the cells with large granules (upper) or nucleolar caps (lower) were observed. Scale bar, 10 μm. (*B* and *C*) Quantification of longest axis (*B*) and area (*C*) of the FCs in cells under indicated conditions. Each scatter dot plot shows the mean (black line). Dots indicate all points of quantified data ($n$ = 500). Mean longest axes of the FCs is shown below: 0 μM: 0.5196 μm, 0.25 μM: 0.6367 μm, 0.5 μM: 0.7192 μm, 1 μM: 1.059 μm. Mean areas of the FCs are shown below: 0 μM: 0.1155 $\mu m^2$, 0. 25 μM: 0.2171 $\mu m^2$, 0.5 μM: 0.3058 $\mu m^2$, 1 μM: 0.6544 $\mu m^2$. Statistical analyses using Kruskal-Wallis test with Dunn's multiple comparison test were performed and the results are shown as follows. (*B*) 0 μM vs 0.25 μM: $P < 0.0001$, 0 μM vs 0.5 μM: $P < 0.0001$, 0 μM vs 1 μM: $P < 0.0001$, 0.25 μM vs 0.5 μM: $P = 0.0211$, 0.25 μM vs 1 μM: $P < 0.0001$, 0.5 μM vs 1 μM: $P < 0.0001$. (*C*) 0 μM vs 0.25 μM: $P < 0.0001$, 0 μM vs 0.5 μM: $P < 0.0001$, 0 μM vs 1 μM: $P < 0.0001$, 0.25 μM vs 0.5 μM: $P = 0.0012$, 0.25 μM vs 1 μM: $P < 0.0001$, 0.5 μM vs 1 μM: $P < 0.0001$. (*D* and *E*) Graphs showing the mean longest axis (*D*) and area (*E*) of the FCs with SEM vs pre-rRNA expression levels. The pre-rRNA expression level in untreated cells is defined as 1.

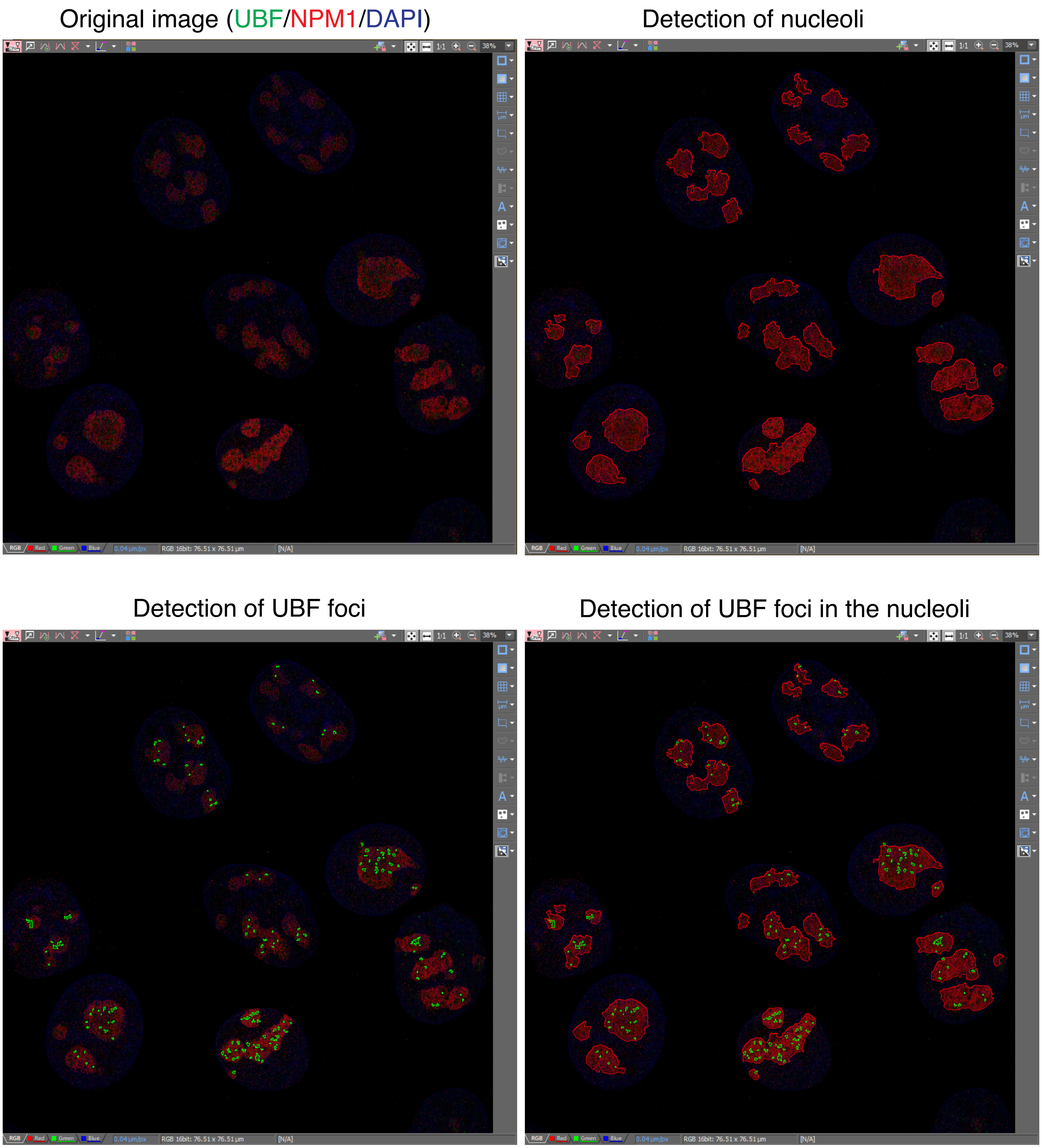


**Figure S4 Quantification of the FCs in the nucleoli**
Screenshots of detections of FCs and GCs by NIS Elements Advanced Research software (NIKON) are shown. Green circles indicate UBF foci, and red circles indicate nucleoli labeled by NPM1 antibodies. UBF foci detected in the nucleoli (lower right) were quantified.
